# Regulation and impact of cardiac lymphangiogenesis in pressure-overload-induced heart failure

**DOI:** 10.1101/2021.04.27.441616

**Authors:** C Heron, A Dumesnil, M Houssari, S Renet, A Lebon, D Godefroy, D Schapman, O Henri, G Riou, L Nicol, JP Henry, M Pieronne-Deperrois, A Ouvrard-Pascaud, R Hägerling, H Chiavelli, JB Michel, P Mulder, S Fraineau, V Richard, V Tardif, E Brakenhielm

## Abstract

**Rationale:** Lymphatics are essential for cardiac health, and insufficient lymphatic expansion (lymphangiogenesis) contributes to development of heart failure (HF) after myocardial infarction. However, the regulation and impact of lymphatics in non-ischemic cardiomyopathy induced by pressure-overload remains to be determined.

**Objective:** Investigate cardiac lymphangiogenesis following transverse aortic constriction (TAC) in adult male or female C57Bl/6J or Balb/c mice, and in patients with end-stage HF.

**Methods & Results:** Cardiac function was evaluated by echocardiography, and cardiac hypertrophy, lymphatics, inflammation, edema, and fibrosis by immunohistochemistry, flow cytometry, microgravimetry, and gene expression analysis, respectively. Treatment with neutralizing anti-VEGFR3 antibodies was applied to inhibit cardiac lymphangiogenesis in mice.

The gender- and strain-dependent mouse cardiac hypertrophic response to TAC, especially increased ventricular wall stress, led to lymphatic expansion in the heart. Our experimental findings that ventricular dilation triggered cardiac lymphangiogenesis was mirrored by observations in clinical HF samples, with increased lymphatic density found in patients with dilated cardiomyopathy. Surprisingly, the striking lymphangiogenesis observed post-TAC in Balb/c mice, linked to increased cardiac Vegfc, did not suffice to resolve myocardial edema, and animals progressed to dilated cardiomyopathy and HF. Conversely, selective inhibition of the essentially Vegfd-driven capillary lymphangiogenesis observed post-TAC in male C57Bl/6J mice did not significantly aggravate cardiac edema. However, cardiac immune cell levels were increased, notably myeloid cells at 3 weeks and T lymphocytes at 8 weeks. Moreover, while the TAC-triggered development of interstitial cardiac fibrosis was unaffected by anti-VEGFR3, inhibition of lymphangiogenesis increased perivascular fibrosis and accelerated the development of left ventricular dilation and cardiac dysfunction.

**Conclusions:** We demonstrate for the first time that endogenous cardiac lymphangiogenesis limits pressure-overload-induced cardiac inflammation and perivascular fibrosis, thus delaying HF development. While these findings remain to be confirmed in a larger study of HF patients, we propose that under settings of pressure-overload poor cardiac lymphangiogenesis may accelerate HF development.

## Introduction

Cardiac lymphatics are essential for maintenance of cardiac health by ensuring tissue fluid homeostasis through the absorption and return of excess extracellular fluid and solutes, but also metabolic waste products, from the heart back to the blood circulation ^1^. In addition, the cardiac lymphatic vasculature, similar as in other organs, regulates the immune response to injury. Indeed, cardiac lymphatics have the potential to modulate immune responses both locally in the myocardium, by actively recruiting and evacuating cardiac-infiltrating immune cells, and distally, at the level of cardiac-draining lymph nodes (dLN), by mediating the uptake from the myocardial interstitium of cytokines, auto-antigens, or pathogens for delivery to dLNs. In addition, recent research has revealed that lymphatic endothelial cells may act directly as antigen-presenting cells, potentially favoring tolerance to self ^2^.

Cardiac inflammation is a current therapeutic target in cardiovascular diseases, notably acute and chronic heart failure (HF). Whereas the contribution of the innate immune system to cardiovascular remodeling, such as after myocardial infarction (MI), has been increasingly recognized ^3^, the role of the adaptive immune system, both B and T lymphocytes, in HF development has been less investigated. Intriguingly, recent experimental and clinical findings indicate that MI and HF are associated with signs of auto-immune disease, including development of autoreactive T cells and anti-cardiac auto-antibodies ^4,5^.

In this context, therapeutic lymphangiogenesis emerges as a novel modality to limit both chronic edema and inflammation in the heart post-MI to prevent HF development ^6–8^. Recently, these experimental findings were extended to models of non-ischemic hypertensive heart disease, induced by either salt-loading in rats or chronic angiotensin-II infusion in mice ^9,10^. Promisingly, lymphangiogenic therapy with Vegfc limited cardiac hypertrophy and remodeling in these settings. However, as the treatment also prevented kidney dysfunction and reduced chronic hypertension ^10,11^ it still remains unknown how much of the functional cardiac benefit observed was directly due to expansion of lymphatics in the heart. In particular, the question remains whether insufficient cardiac lymphangiogenesis may contribute to the chronic myocardial edema, inflammation, fibrosis, and cardiac remodeling that occurs during severe cardiac hypertrophy in response to pressure-overload, such as in patients with aortic stenosis. We hypothesized that increased cardiac wall stress during pressure-overload may induce Vegfc and/or Vegfd growth factors to stimulate cardiac lymphangiogenesis, similar as previously reported for Vegfa and cardiac angiogenesis during physiological cardiac growth ^12,13^. Further, we postulated that insufficient lymphangiogenesis may lead to accelerated HF development and cardiac decompensation (reduced cardiac output). Of note, insufficient angiogenesis contributes to cardiac decompensation in pathological hypertrophy ^14^. Here, we examined the regulation and impact of lymphangiogenesis in the heart during the switch from compensated to decompensated cardiac hypertrophy using mouse models of pressure-overload induced by transversal aortic constriction (TAC).

## Materials & Methods

Cardiac lymphatic remodeling was investigated in adult male and female C57Bl/6 mice and female Balb/c mice (Janvier Laboratories, France) following TAC. Briefly, a minimally invasive method^15^ was used to constrict the aortic arch, using a 26G needle. Double-banding of the aorta was applied to prevent suture internalization and model variability, as described^16^. Modulation of cardiac lymphangiogenesis was performed using a rat anti-mouse VEGFR3 blocking antibody, mF4-31C1 (Imclone/Eli Lilly), as described ^17^. The antibody therapy consisted of repeated (starting from day 7 post-TAC) twice a week i.p injections of 800 µg/mouse of mF4-31C1 or rat IgG as a control. Cardiac samples from end-stage HF patients (recipients of cardiac transplants at the Bichat Hospital in Paris, France) were examined by immunohistology. Discarded cardiac autopsy samples, obtained from donors without cardiovascular disease, were used as healthy controls. For details see *Suppl. Methods*.

### Cardiac functional, cellular and molecular analyses in mice

Cardiac function was evaluated by echocardiography ^8^. Cardiac sections were analyzed by histology and immunohistochemistry to determine lymphatic and blood vessel densities and sizes, immune cell infiltration, cardiomyocyte hypertrophy and fibrosis. Whole mount-staining of cardiac lymphatics and arterioles was performed followed by a modified iDISCO+ clearing protocol for imaging by light sheet (ultramicroscope II, LaVision BioTec) and confocal laser scanning (Leica SP8) microscopy ^8^. Cardiac gene expression was analyzed by RT-qPCR ^8^. For details see *Suppl. Methods*.

### Flow-cytometry

Cardiac immune cells were analyzed by flow cytometry using a LSRFortessa (BD Biosciences) cytometer ^8^. Results are expressed cells per mg cardiac tissue. For details see *Suppl. Methods*.

### Study approval

Anonymized human heart samples evaluated in this study were obtained with informed consent by the Bichat hospital biobank (*U1148 BB-0033-00029/* BBMRI, coordinator JB Michel) authorized for tissue collection by the Inserm institutional review board. Animal experiments performed in this study were submitted to the regional Normandy ethics review board in line with E.U and French legislation (01181.01 / APAFIS #8157-2016121311094625 v5 and APAFIS #23175-2019112214599474 v6). A total of 224 C57Bl/6 male or female mice and 58 Balb/c female mice, surviving TAC or sham-operation were included in this study.

### Statistics

Data are presented as mean ± s.e.m. Comparisons were selected to determine: 1) impact of pathology (healthy sham *vs*. TAC); 2) effect of treatment (anti-VEGFR3-treated vs. IgG TAC controls). Statistical analyses for comparisons of two independent groups were performed using either Student’s two-tailed t-test for groups with normal distribution, or alternatively by Mann Whitney U test for samples where normality could not be ascertained based on D’Agostino & Pearson omnibus normality test. For comparisons of three groups or more either One-way ANOVA followed by Bonferroni posthoc (for parameters with n>7 with normal distribution), or alternatively Kruskal-Wallis nonparametric analysis followed by Dunn’s posthoc multiple comparison (for parameters with non-Gaussian distribution) were performed. Longitudinal echocardiography studies were analyzed by paired two-way ANOVA followed by Bonferroni posthoc, while morphometric data and gene expression data were analyzed by two-way ANOVA followed by Sidak’s posthoc for pair-wise comparisons or Dunnett’s posthoc to compare 3 groups. Non-parametric Spearman rank order tests were used for evaluating correlations. All analyses were performed using GraphPad Prism software.

## Results

### Strain- and gender-dependent cardiac response to pressure-overload

The transverse aortic constriction (TAC) rapidly led to severe cardiac hypertrophy in both male and female C57Bl/6J mice, with a 50% increase in left ventricular mass observed by 3 weeks (**Suppl Fig. 1a-b**). At 8 weeks post-TAC, the cardiac hypertrophy in C57Bl/6J mice had stabilized, with a 30% increase in cardiomyocyte sizes as compared to sham-operated controls (**Fig. 1a-d**). In contrast, female Balb/c displayed a weaker initial hypertrophic response, despite a similar increase post-TAC in transaortic pressure gradients in both strains, and only by 8 weeks was the cardiac mass and cardiomyocyte hypertrophy significantly increased (**Fig. 1a-d, Suppl Fig. 1a-b**). At the end of the study, the degree of cardiac hypertrophy induced by the TAC procedure was comparable between strains, with more concentric hypertrophy observed in female C57Bl/6J and more eccentric hypertrophy in the other TAC groups. Conversely, while body weight gain was unaffected by TAC in C57Bl/6J mice, it was slightly reduced in Balb/c mice by 8 weeks (**Suppl. Fig. 1c**). Interestingly, the severe cardiac hypertrophy led to left ventricular dilation and cardiac decompensation only in male C57Bl/6J and female Balb/c, but not in female C57Bl/6J mice. Indeed, while both male C57Bl/6J and female Balb/c mice progressed to HF by 6-8 weeks post-TAC, C57Bl/6J females had not yet developed overt cardiac dysfunction by the end of the study (**Suppl. tables 1-3**). Analyses of cardiac expression of ANP and BNP genes (*Nppa* and *Nppb*, respectively), used as indicators of end-diastolic wall stress (EDWS) following pressure-overload^18,19^, revealed an earlier and more robust increase in male C57Bl/6J post-TAC as compared to females (**Suppl. Fig. 1d**). However, at 8 weeks, *Nppa* levels were comparable in male C57Bl/6J and female Balb/c mice (**Fig. 1e**). The angiogenic response was coordinated with the degree of cardiac hypertrophy, with the largest increase in blood vessel to cardiomyocyte ratios observed in male C57Bl/6J at 8 weeks post-TAC, with a smaller, non-significant increase in female Balb/c (**Fig. 1f, g**).

**Fig. 1.**
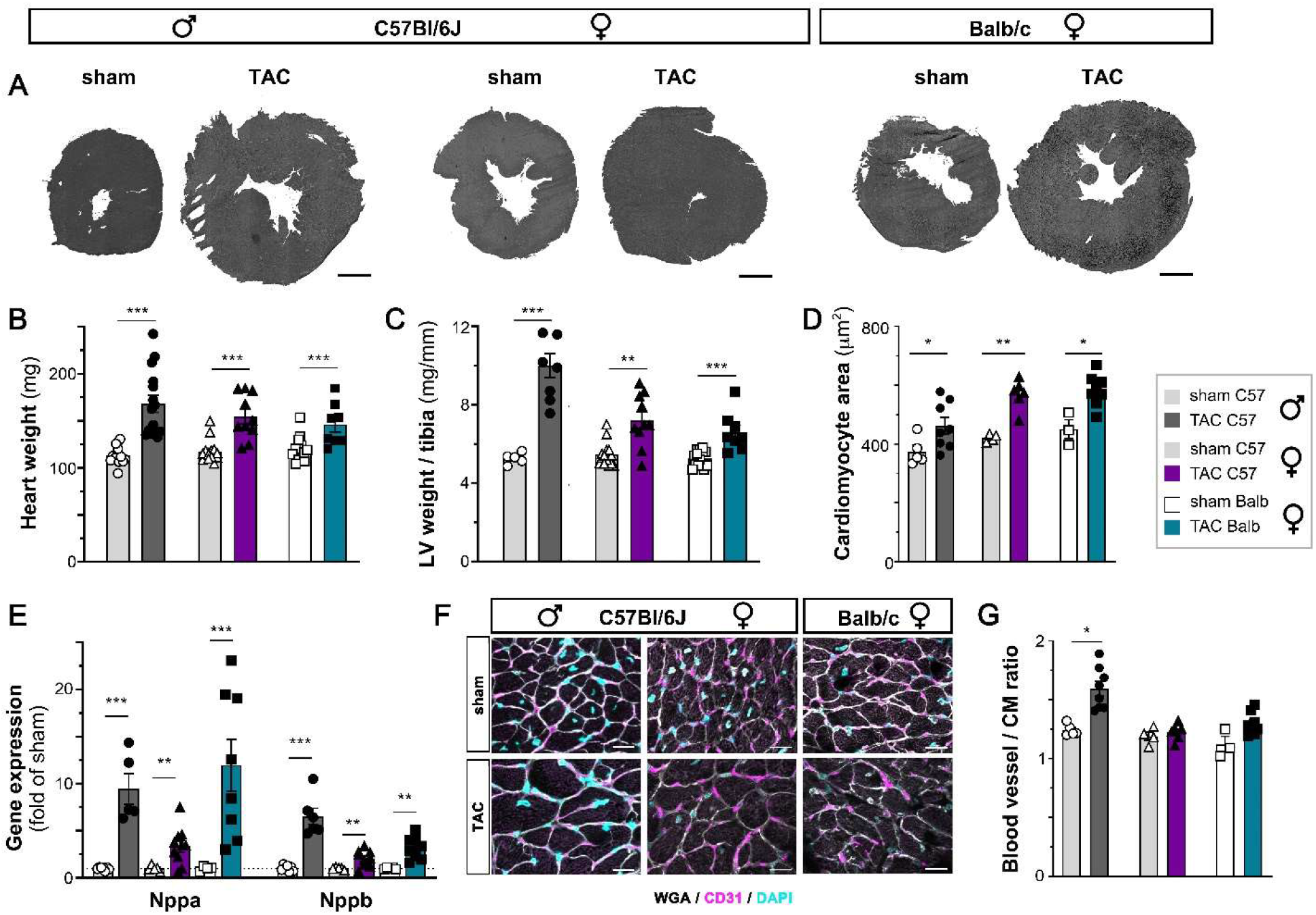
Evaluation of cardiac hypertrophy at 8 weeks post-TAC. Examples (**a**, scale bar 1 mm) and morphometric assessment of heart weights (**b**), and left ventricular (LV) weights normalized to tibia lengths (**c**) at 8 weeks in male C57Bl/6J mice: sham (open circles, *n=13*) or TAC (closed circles, *dark grey bar n=15*); female C57Bl/6J mice: sham (open triangles, *n=14*) or TAC (closed triangles, *purple bar n=10*); and in Balb/c female mice: sham (open square, *n=14*) or TAC (closed square, *blue bar n=8*). Analysis of cardiomyocyte cross-sectional area at 8 weeks (**d**, *n=3-8* animals per group). Cardiac expression analyses of *Nppa* and *Nppb* (**e**) at 8 weeks (*n=5-10* per group). Examples (**f**) and evaluation (**g**) of blood vessel to cardiomyocyte ratios at 8 weeks post-TAC (*n=3-8* animals per group). WGA, *white*, CD31, *purple*, Dapi, *blue* (scale bar 20 µm). Groups were compared pair-wise by two-way ANOVA followed by Sidak’s multiple comparison test (for morphometry and cardiac expression analyses) or by non-parametric Kruskal Wallis followed by Dunn’s posthoc test (for immunohistochemistry) * p<0.05, ** p<0.01, *** p<0.001 *versus* sham.

### Regulation of lymphatic remodeling during cardiac hypertrophy

Next, we investigated cardiac lymphatic remodeling. We hypothesized that cardiac lymphangiogenesis would accompany the cardiac hypertrophy to match the drainage capacity to the expanded ventricular mass. We thus expected an earlier and more robust cardiac lymphatic expansion in C57Bl/6J as compared to Balb/c mice. However, both immunohistochemistry and cardiac gene expression analyses revealed that at 3 weeks post-TAC there was no significant increase in lymphatic density, nor in cardiac *Vegfc* or *Vegfd* gene expression, in C57Bl/6 males despite their extensive cardiac hypertrophy (**Fig. 2b, Suppl. Fig 2a**). Further, although expression of the lymphatic chemokine *Ccl21* was increased (**Suppl. Fig 2b**), indicating lymphatic activation, the open lymphatic density in the heart was not significantly increased (**Fig. 2c**). However, cardiac precollector slimming, previously reported post-MI in rodents ^6,8^, did not occur, and subsequently open cardiac lymphatic area remained unchanged (**Suppl. Fig. 2 d**). Only by 8 weeks post-TAC was the cardiac expression of Vegfc, and especially Vegfd, significantly increased in C57Bl/6J males (**Fig. 2d**). Consequently, the expression of lymphatic markers podoplanin (*Pdpn*) and *Ccl21* was increased (**Fig. 2e**), together with slightly increased capillary lymphatic density at 8 weeks (**Fig. 2b**). Thus, the rapid and severe cardiac growth induced by TAC in C57Bl/6J males was associated with a slow and moderate lymphangiogenic response, essentially driven by Vegfd. Surprisingly, different from males, at 3 weeks post-TAC the cardiac hypertrophy in C57Bl/6J females was associated with a transient increase in both cardiac lymphatic density (**Fig. 2b**) and open lymphatic area (**Suppl. Fig. 2d**). In agreement with early cardiac lymphatic expansion, both *Pdpn* and *Ccl21* cardiac gene expression levels were increased (**Suppl. Fig 2b**). However, by 8 weeks post-TAC, the lymphatic density, as well as cardiac expression of lymphatic markers, had essentially reverted back to initial levels (**Fig. 2a-e**). Thus, despite similar cardiac hypertrophy post-TAC as in males, female C57Bl/6J mice displayed only transient, likely Vegfd-driven, cardiac lymphangiogenesis. Finally, in Balb/c mice, where the same pressure-overload stimuli induced a slower hypertrophic response, there was no cardiac lymphangiogenesis at 3 weeks post-TAC (**Fig. 2b, c; Suppl. Fig. 2a, b, d**). Interestingly, the dilated cardiac phenotype observed at 6-8 weeks in female Balb/c mice (**Suppl. table 2**), characterized by wall thinning indicative of severe and chronically elevated cardiac wall stress, was associated with a strong and selective increase in cardiac *Vegfc* gene expression (**Fig. 2d**). This led to a remarkably potent cardiac lymphangiogenic response by 8 weeks post-TAC (**Fig. 2b, c, e**), with expansion mostly of lymphatic capillaries, as observed by whole mount imaging (**Fig. 2a**).

**Fig. 2.**
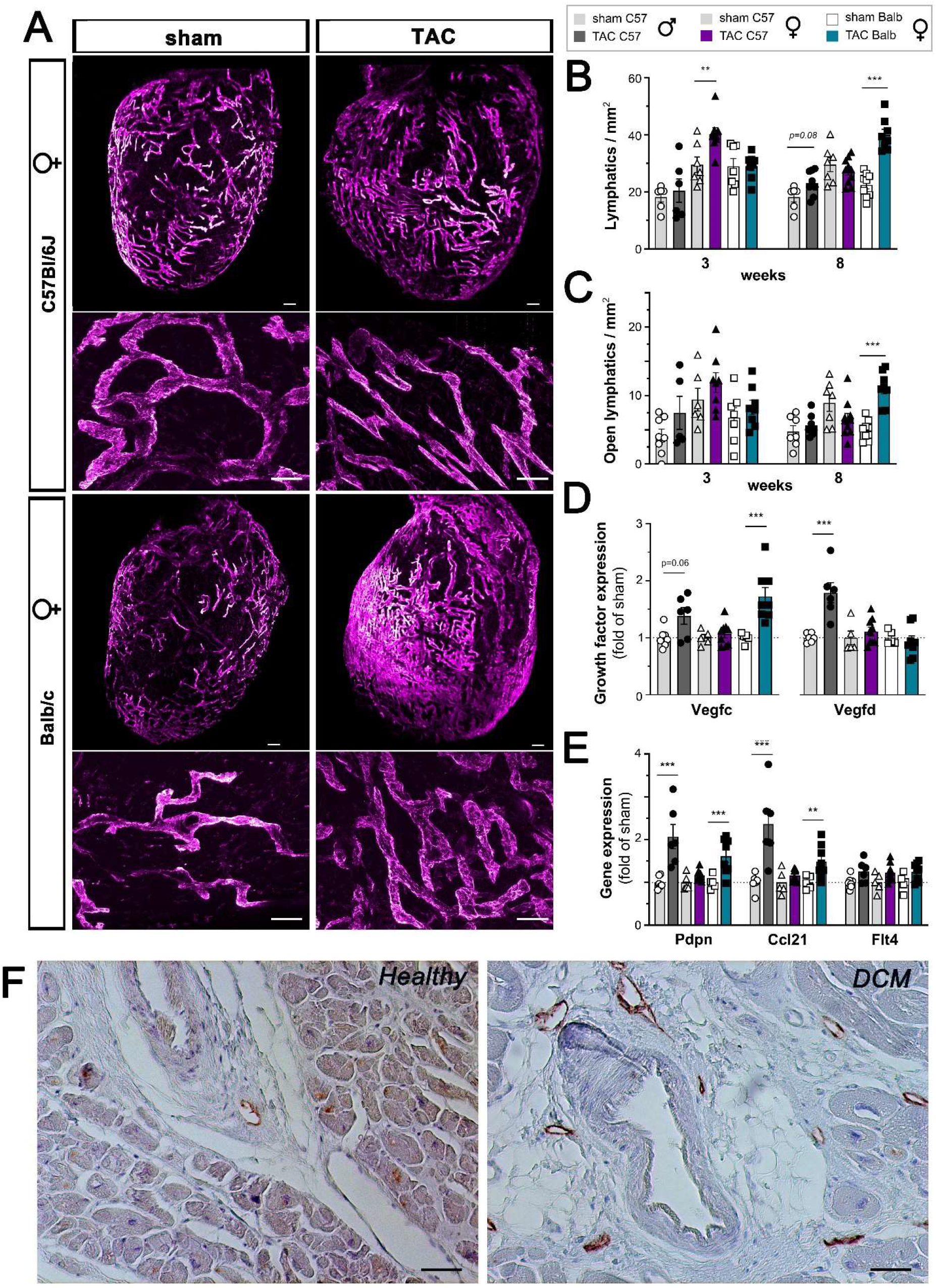
Evaluation of cardiac lymphatic remodeling post-TAC. Examples by light sheet and confocal imaging of cardiac lymphatics (**a**) at 8 weeks post-TAC in female C57Bl/6J or Balb/c mice. Lyve1, *purple*. (Scale bar *upper row* 300 µm, *lower row* zoomed confocal views 100 µm). Evaluation of total lymphatic vessel density (**b**) and open lymphatic density (**c**) in the LV subepicardium at 3 or 8 weeks in C57Bl/6J males: sham (open circles, *n=6*) or TAC (closed circles, *dark grey bar n=8*); C57Bl/6J females: sham (open triangles, *n=7*) or TAC (closed triangles, *purple bar n=10*); and in Balb/c females: sham (open square, *n=9*) or TAC (closed square, *blue bar n=8*). Cardiac expression analyses (*n=5-10* per group) at 8 weeks of *Vegfc* and *Vegfd* (**d**), and lymphatic markers *Pdpn, Ccl21*, and *Flt4* (**e**). Groups were compared pair-wise by non-parametric Kruskal Wallis followed by Dunn’s posthoc test (for immunohistochemistry) or Sidak’s multiple comparisons test (for cardiac expression analyses) * p<0.05, ** p<0.01, *** p<0.001 *versus* sham. Examples of septal cardiac lymphatic density (**f**) in a healthy cardiac donor versus an end-stage HF patient with dilated cardiomyopathy (DCM). Podoplanin, *brown*. (Scale bar 50 µm).

Taken together, we found that during pressure-overload both *Vegfc* and *Vegfd* cardiac gene expression positively correlated with augmented EDWS (approximated by increased cardiac *Nppa* and *Nppb* levels) (**Suppl. Fig. 3a, b**), rather than with the degree of cardiac hypertrophy (increased left ventricular mass). Similarly, in humans, we found that the lymphatic density, notably of perivascular lymphatics, in the interventricular septum was most increased in HF patients with primitive dilated cardiomyopathy (DCM), as compared to healthy controls (**Fig. 2f, table I**).

**Table I.** Immunohistochemical analyses of lymphatics in the cardiac septum of HF patients and healthy controls. *ICM*, ischemic cardiomyopathy; *DCM*, dilated cardiomyopathy–

|  | *n* | Age<br>(years) | Male<br>gender<br>(%) | total<br>lymphatics<br>/ mm<sup>2</sup> | open<br>lymphatics<br>/ mm<sup>2</sup> | lymphatic<br>diameter<br>(μm) | perivascular<br>lymphatics /<br>mm<sup>2</sup> |
| --- | --- | --- | --- | --- | --- | --- | --- |
| Healthy controls | 4 | 53 ± 5 | 100% | 12 ± 4 | 10 ± 3 | 27 ± 10 | 4 ± 2 |
| ICM | 6 | 57 ± 6 | 100% | 16 ± 2 | 13 ± 2 | 30 ± 2 | 6 ± 1 |
| Primitive DCM | 3 | 50 ± 2 | 67% | 19 ± 3 | 17 ± 2 | 23 ± 3 | 12 ± 7 |

ICM, ischemic cardiomyopathy; DCM, dilated cardiomyopathy–

In our mouse model, the lymphangiogenic response seemed to be influenced by gender- and strain-dependent differences. One such inherent strain-dependent difference is the cytokine environment, with a significantly higher cardiac expression observed after TAC in Balb/c mice of the pro-inflammatory cytokines interleukin (IL)-1β and IL-6, as compared with in C57Bl/6J mice (**Fig. 3a**). Of note, previous work has shown that both these cytokines induce Vegfc production in fibroblasts and macrophages to stimulate lymphangiogenesis in other organs ^20–22^. Thus, one explanation for the more potent lymphatic response to pressure-overload in Balb/c mice may be the higher cardiac IL-1β and IL-6 levels that, together with elevated wall stress due to left ventricular dilation, drive cardiac *Vegfc* expression. Indeed, we found that cardiac *IL-1β* levels, better than *IL-6*, positively correlated with *Vegfc*, but not with *Vegfd* or *Vegfa*, gene expression in the heart (**Suppl. Fig 3c**). Concerning gender-dependent differences in C57Bl/6J mice, estrogen has been shown to increase, *via* estrogen receptor-α, the expression of Vegfd, as well as Vegfr3 ^23^. However, we did not observe any effects of gender on cardiac *Vegfd* nor Vegfr3 (*Flt4*) gene expression, neither at baseline or after pressure-overload (**Fig. 2d, e**).

**Fig. 3.**
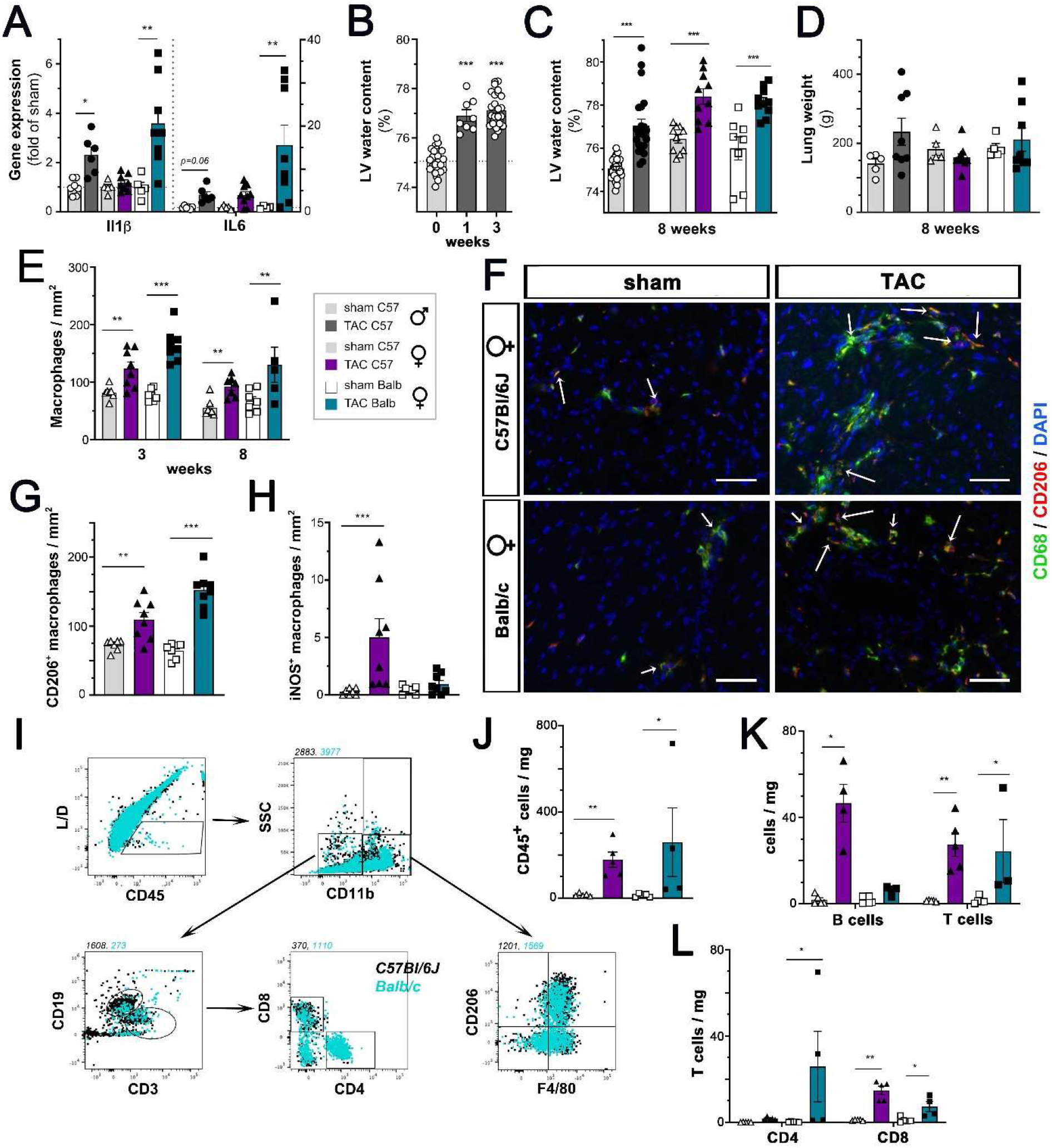
Kinetics of edema and inflammation after TAC. Cardiac expression analysis of *Il1β* and *Il6* (**a**) at 8 weeks (*n=5-10* animals per group). Assessment of myocardial water content at 1 and 3 weeks post-TAC in male C57Bl/6J mice (**b**) or at 8 weeks (**c**) in C57Bl/6J males: sham (open circles, *n=8*) or TAC (closed circles, *dark grey bar n=8*); C57Bl/6J females: sham (open triangles, *n=8*) or TAC (closed triangles, *purple bar n=10*); and in Balb/c females: sham (open square, *n=8*) or TAC (closed square, *blue bar n=10*). Evaluation of pulmonary weights at 8 weeks (**d**). Quantification of cardiac-infiltrating macrophage density (**e**) at 3 or 8 weeks in C57Bl/6J females: sham (*n=7*) or TAC (*n=8*); and in Balb/c females: sham (*n=7*) or TAC (*n=8*). Examples of cardiac macrophages (**f**) at 3 weeks. CD68, *green*, CD206, *red*; dapi *blue*. Arrows indicate CD206^+^ macrophages (scalebar, 50 µm). Quantification of non-classical CD206^+^ macrophages (**g**) and classical iNOS^+^ macrophages (**h**) at 3 weeks post-TAC. Examples of flow cytometry gating in TAC mice (**i**). Quantification in C57Bl/6J and Balb/c females at 8 weeks post-TAC (*n=4-5 samples* per group) of cardiac-infiltrating CD45^+^ immune cells (**j**), CD19^+^ B cells and CD3^+^ T cells (**k**), and CD4^+^ vs CD8^+^ T cell subpopulations (**l**). Groups were compared pair-wise by two-way ANOVA followed by Sidak’s multiple comparison tests (morphometry data and expression analyses) or non-parametric Kruskal Wallis followed Dunn’s posthoc test (for microgravimetry and immunohistochemistry) or by Mann Whitney U test (for flow cytometry); * p<0.05, ** p<0.01, *** p<0.001 *versus* sham.

### Cardiac lymphangiogenesis does not resolve myocardial edema after TAC

Next, we set out to investigate the functional effects linked to cardiac lymphatic remodeling during pathological cardiac hypertrophy. First, we observed that severe myocardial edema developed rapidly after pressure-overload (**Fig. 3b**). Surprisingly, there was no difference in the degree of cardiac edema at 8 weeks between C57Bl/6J and Balb/c mice (**Fig. 3c**), indicating that the prominent cardiac lymphangiogenesis occurring in the latter was not sufficient to resolve the chronic edema. In contrast, pulmonary edema occurred by 3 weeks post-TAC in C57Bl/6J male mice (**Suppl. Fig. 4a**), and only tended to increase in Balb/c females at 8 weeks (**Fig. 3d)**. This development coincided with the appearance of left ventricular dilation and a drop in cardiac output, as observed by echocardiography at 3 weeks post-TAC in C57Bl/6J male mice and at 6-8 weeks in Balb/c females (**Suppl. tables 1, 2**). In contrast, female C57Bl/6J mice, which did not progress to decompensated cardiac hypertrophy during the 8-week study (**Suppl. table 3**), displayed pulmonary weights similar to healthy sham controls.

### Impact of lymphangiogenesis on cardiac immune cell levels

Another aspect of lymphatic function is the clearance of immune cells to resolve inflammation. We and others previously demonstrated that therapeutic lymphangiogenesis limits cardiac inflammation post-MI in rats and mice ^6–8^. Conversely, prior studies have elegantly demonstrated the key role that different immune cells play during the transition from physiological to pathological hypertrophy following pressure-overload ^24–27^. We thus set out to investigate whether the graded cardiac lymphangiogenic response to cardiac hypertrophy observed in our models was associated with differential clearance of immune cells. Based on our lymphatics data, we expected better resolution of cardiac inflammation at 8 weeks in Balb/c versus C57Bl/6J female mice. First, in agreement with the literature, we found, using immunohistochemistry and flow cytometry, that the early immune response to pressure-overload was characterized by increased cardiac immune cell levels, notably monocytes and alternative CD206^+^ macrophages (**Fig. 3e, f, g, Suppl. Fig. 4b, c, e**). Cardiac macrophages may be a rich source of Vegfc, as indicated by our immunohistochemical analyses at 3 weeks post-TAC (**Suppl. Fig. 4g**). Interestingly, the level of classical, iNOS^+^ pro-inflammatory macrophages was highest in C57Bl/6J females, which displayed concentric hypertrophy and waning lymphangiogenesis (**Fig. 3h**). At 8 weeks post-TAC, both C57Bl/6J and Balb/c females exhibited increased T cell infiltration (**Fig. 3i-k**). Interestingly, while in Balb/c this included expansion of both CD4^+^ and CD8^+^ T cells, in C57Bl/6J mice only CD8^+^ T cell levels were increased (**Fig. 3i, l**). Conversely, the cardiac level of mature natural killer (NK) cells was significantly increased post-TAC only in female C57Bl/6J mice (**Suppl. Fig. 4d**). Further, only in C57Bl/6J female mice was there an increase in cardiac B cell levels (**Fig. 3k**) and *TNFα* expression (**Suppl. Fig. 4f**) at 8 weeks post-TAC. However, this cardiac immune response was not sufficient to induce overt cardiac dysfunction in C57Bl/6J females, as noted above. Of note, while some studies indicate a key pathological role of B lymphocytes to promote ventricular dilation following pressure-overload ^27^, these cells have also been suggested to serve a beneficial role post-TAC by producing immune-suppressive IL-10 to signal inflammatory resolution ^26^. These different properties may be carried by distinct B cells subsets, as recently suggested by a cardiac scRNASeq study post-TAC ^28^. Taken together, our data suggest that a poor lymphangiogenic response in the heart during pathological cardiac hypertrophy may contribute to an increase in pro-inflammatory cardiac immune cells levels, notably iNOS^+^ classical macrophages and B lymphocytes.

### Inhibition of lymphangiogenesis aggravates cardiac hypertrophy and accelerates cardiac dysfunction

To further test the hypothesis that even a moderate lymphangiogenic response, as observed in male C57Bl/6J mice, may have protected against cardiac inflammation and/or decompensation after pressure-overload, we next investigated the impact of selective inhibition of lymphangiogenesis using a VEGFR3 blocking antibody, mF4-31C1, as previously described ^17^. To not interfere with the initial response to the pressure-overload, administration of the blocking antibody, or non-specific rat IgG in controls, was initiated at 1 week after TAC surgery, and the treatment was then maintained through the 8-week study. We found that the anti-VEGFR3 treatment rapidly, selectively, and completely blocked cardiac lymphangiogenesis post-TAC, despite persistently elevated cardiac *Vegfd* expression (**Fig. 4a-d; Suppl. Fig. 5a, b**). Indeed, while the treatment prevented the upregulation of cardiac *Ccl21* expression already at 3 weeks post-TAC (**Suppl. Fig. 5c**), by 8 weeks all investigated lymphatic markers (*Pdpn, Ccl21, Flt4*) were reduced as compared to TAC controls (**Fig. 4e**). The inhibition of lymphangiogenesis affected both lymphatic capillaries and precollectors, as demonstrated by cardiac whole mount imaging (**Fig. 4a**). 3D analyses further revealed a reduction of both lymphatic volume density and the frequency of short vessel segments, indicating prevention of lymphatic sprouting post-TAC by anti-VEGFR3 (**Fig. 4f-i**). In contrast, the treatment had no effect on cardiac angiogenesis or arteriogenesis, nor on cardiac perfusion, as evaluated by MRI (**Suppl. Fig 6**).

**Fig. 4.**
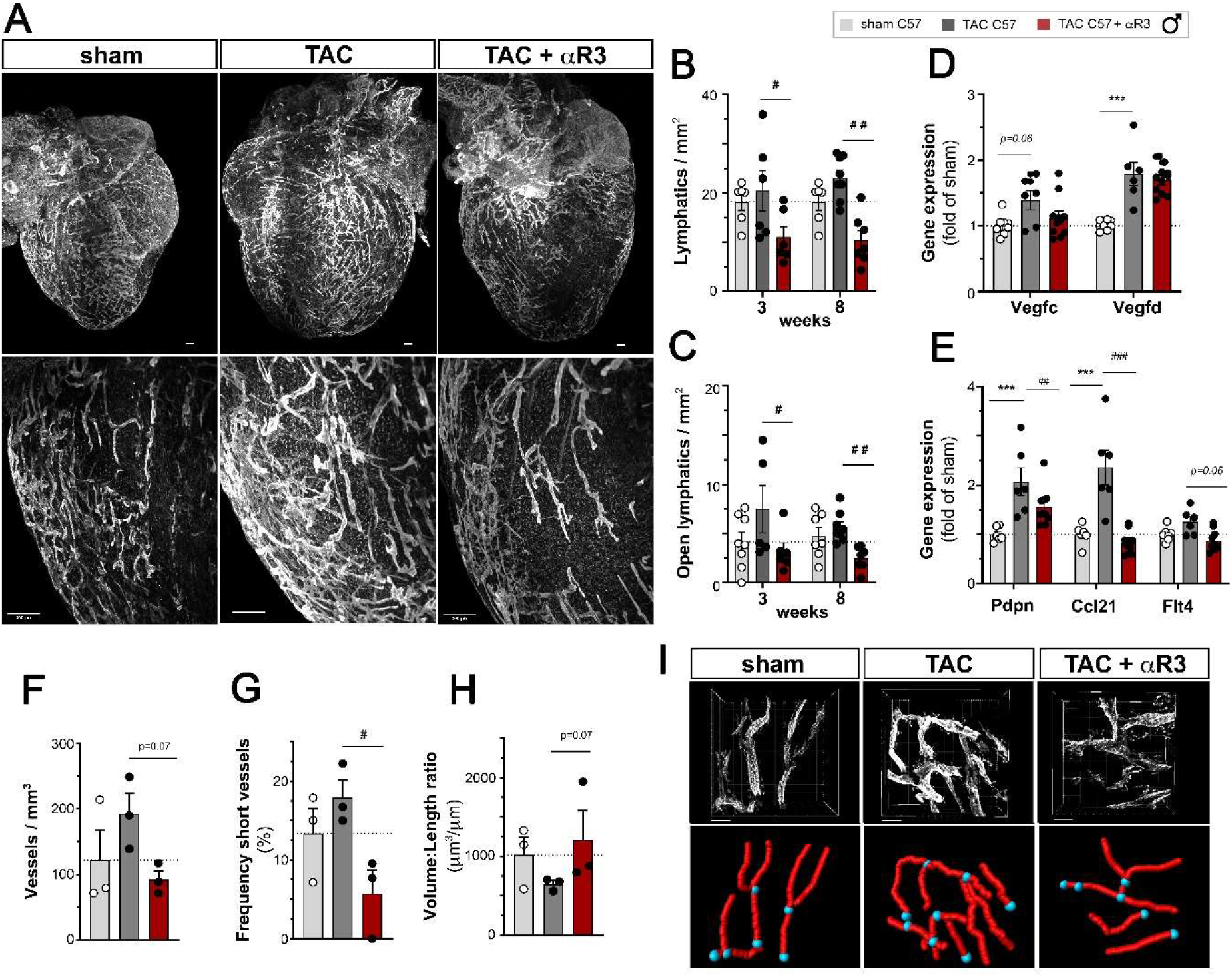
Anti-VEGFR3 treatment blocks cardiac lymphangiogenesis post-TAC. Light sheet imaging of cardiac lymphatics (**a**) at 8 weeks post-TAC visualized by Lyve1 (*grey*). (Scale bar 300 µm). Evaluation of total lymphatic vessel density (**b**) and open lymphatic density (**c**) in the LV subepicardium at 3 and 8 weeks in male C57Bl/6J sham (open circles, *n=6*), TAC controls (closed circles, *dark grey bar n=6-8*), and anti-VEGFR3-treated TAC (closed circles, *red bar n=6-7*). Cardiac expression analyses of *Vegfc* and *Vegfd* (**d**) and lymphatic markers *Pdpn, Ccl21*, and *Flt4* (**e**) at 8 weeks (*n=6-10* animals per group). 3D confocal analyses of cardiac lymphatic volume density (**f**), frequency of short lymphatic branches (**g**), and volume:length ratio (**h**) at 3 weeks post-TAC (*n=3* mice per group). Examples of modeling of cardiac lymphatics visualized by Lyve1 (*grey*) (**i**, Scale bar 100 µm). Groups were compared by non-parametric Kruskal Wallis followed by Dunn’s posthoc test (for immunohistochemistry) and by two-way ANOVA followed by Dunnett’s multiple comparisons test (for expression analyses). * p<0.05 *versus* sham, ## p<0.01, ### p<0.001 *versus* control TAC.

Concerning the cardiac hypertrophic response to the pressure-overload, there was a more pronounced increase in cardiac mass in the anti-VEGFR3-treated group by 8 weeks (**Fig. 5a, Suppl. Fig. 7a**). In contrast, there was no change in body weight gain (**Suppl. Fig. 7b**), and cardiac expression of *Nppa* and *Nppb*, used as biomarkers of EDWS, were similarly increased at 3- and 8-weeks post-TAC in both groups (**Fig. 5b, Suppl. Fig. 7c**). However, in the anti-VEGFR3-treated group the development of left ventricular dilation was accelerated, occurring already by 3 weeks post-TAC, as compared to 6 weeks in controls (**Suppl. table 4**). Nevertheless, at the end of the study, the cardiac dysfunction was similar between the two C57Bl/6J male TAC groups, indicating that inhibition of lymphangiogenesis accelerated the progression, but not the severity, of the cardiac decompensation process.

**Fig. 5.**
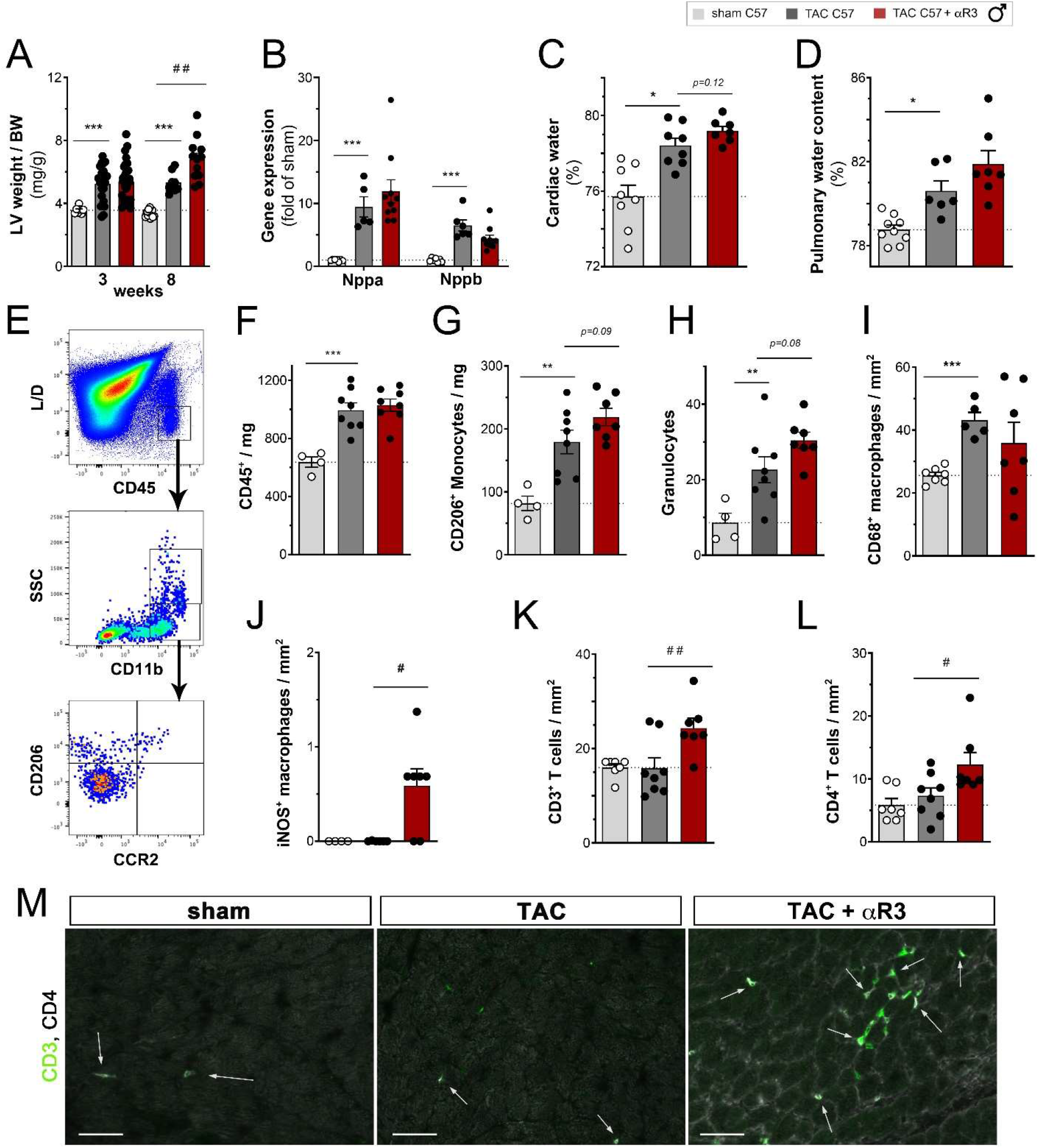
Aggravation of cardiac hypertrophy and inflammation, but not edema, by anti-VEGFR3 treatment. Morphometric assessment of left ventricular (LV) weight normalized to body weight (**a**) at 3 or 8 weeks in male C57Bl/6J sham (open circles, *n=7-13*) or TAC controls (closed circles, *dark grey bar n=8-20*); anti-VEGFR3-treated TAC (closed circles, red bar *n=12-20*). Analysis of cardiac *Nppa* and *Nppb* (**b**) expression at 8 weeks (n=6-10 animals per group). Assessment of myocardial (**c**) and pulmonary (**d**) water content at 3 weeks post-TAC in male C57Bl/6J sham (*n=8-9*), TAC controls (*n=6-8*), and anti-VEGFR3-treated TAC (*n=7*). Flow cytometric evaluation (*n=4-8 per group*) at 3 weeks post-TAC (**e**) of cardiac-infiltrating CD45^+^ immune cells (**f**), CD11b^+^ CD206^+^ alternative monocytes (**g**) and CD11b^+^ SSC^high^ granculocytes (**h**). Data is reported as *cells per mg* cardiac tissue. Quantification by immunohistochemistry at 3 weeks post-TAC (*n=6-7* animals per group) of cardiac-infiltrating CD68^+^ macrophages (**i**), and classical iNOS^+^ macrophages (**j**). Quantification by immunohistochemistry at 8 weeks post-TAC (*n=7-8* animals per group) of cardiac-infiltrating CD3^+^ T cells (**k**), and CD4^+^ T cells (**l**). Examples at 8 weeks post-TAC of cardiac CD3^+^ total T cells (g*reen*) and CD4^+^ T cells (*grey*) indicated by arrows (**m**). Scalebar 50 µm. Groups were compared by non-parametric Kruskal Wallis followed by Dunn’s posthoc test (for immunohistochemistry, microgravimetry and flow-cytometry) and by two-way ANOVA followed by Sidak’s multiple comparison test (for morphometric data) or Dunnett’s multiple comparisons test (for expression analyses). * p<0.05, ** p<0.01, *** p<0.001 *versus* sham, ## p<0.01, ### p<0.001 *versus* control TAC.

### Impact of inhibition of lymphangiogenesis on edema and resolution of cardiac inflammation

To determine by which mechanism inhibition of lymphangiogenesis may have accelerated cardiac hypertrophy and left ventricular dilation, we analyzed cardiac and pulmonary water content at 3 weeks post-TAC. Using microgravimetry, we found that both cardiac and pulmonary edema only tended to be increased in the anti-VEGFR3 treated group as compared to TAC controls (**Fig. 5c, d**). In contrast, using flow cytometry, we found that both granulocytes and monocyte levels in the heart tended to increase at 3 weeks in anti-VEGFR3 treated TAC mice (**Fig. 5e-h, Suppl. Fig 7d**). In contrast, cardiac lymphocyte levels were unchanged as compared to TAC controls (**Suppl. Fig 7e-i**). Further, using immunohistochemistry, we observed that although cardiac macrophage density was not increased, the level of classical pro-inflammatory iNOS^+^ CD206^-^ macrophages was increased in anti-VEGFR3-treated mice (**Fig. 5i, j**), similarly as observed post-TAC in female C57Bl/6J mice. Moreover, by 8 weeks, anti-VEGFR3-treated male mice displayed increased cardiac levels of T cells, especially CD4^+^ helper T cells (**Fig. 5 k-m**), previously demonstrated to accelerate cardiac decompensation ^25^. In contrast, there was no change in cardiac T regulatory cells (**Suppl. Fig. 7g**).

### Links to cardiac fibrosis

Both cardiac edema and inflammation are potent drivers of cardiac fibrosis ^29^. Thus, we next set out to determine whether poor lymphangiogenesis during pressure-overload, by aggravating cardiac inflammation, may have accentuated cardiac collagen production and/or deposition. First, we examined cardiac fibrosis at 8 weeks post-TAC in C57Bl/6J and Balb/c females using Sirius red histological evaluations. Despite their clear differences in cardiac lymphangiogenesis following pressure-overload, the development of interstitial fibrosis post-TAC was found to be similar in both strains (**Fig. 6a, Suppl. Fig 8c**). This apparent absence of impact of lymphatics on development of interstitial fibrosis may reflect our observation that cardiac edema was equally severe in both female TAC groups. However, it should be noted that during pressure-overload, Balb/c females displayed much higher cardiac *Il-6* and *Il-1*β expression (**Fig. 3a**), expected to promote cardiac fibroblast activation. In contrast, perivascular fibrosis was significantly increased only in C57Bl/6J females, characterized by poor lymphangiogenesis and increased cardiac pro-inflammatory cell infiltration (**Fig. 6b, d**).

**Fig. 6.**
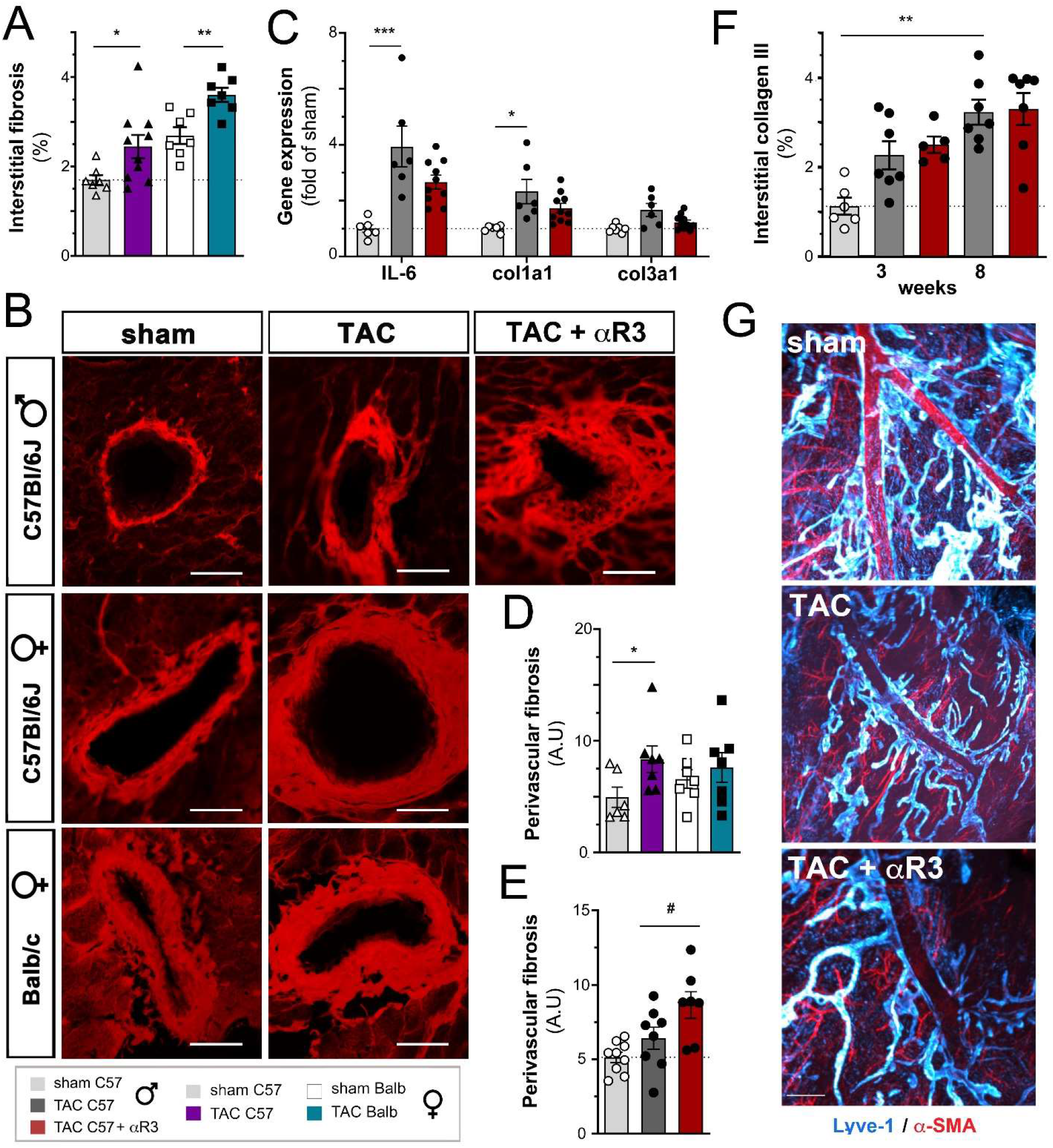
Cardiac interstitial and perivascular fibrosis and perivascular lymphangiogenesis post-TAC. Quantification of interstitial fibrosis (**a**) at 8 weeks in female C57Bl/6J sham (open triangles, *n=7*) or TAC (closed triangles, *purple bar n=10*) and in female Balb/c sham (open square, *n=7*) or TAC (closed square, *blue bar n=8*). Examples of perivascular fibrosis (**b**) evaluated as relative fibrotic area surrounding arterioles in the size range of 5-50 µm diameter. (Scalebar 50 µm). Quantification of perivascular fibrosis at 8 weeks in female C57Bl/6J or Balb/c mice (**d**). Cardiac gene expression of *Il6, Col1a1* and *Col3a1* (**c**) analyzed at 8 weeks post-TAC in male C57Bl/6J mice (*n=6-10* per group). Quantification of interstitial collagen III density (**f**) at 3 and 8 weeks in male C57Bl/6J sham (open circles, *n=6-9*), TAC controls (closed circles, *dark grey bar n=5-7*); and anti-VEGFR3-treated TAC (closed circles, *red bar n=5-7*). Quantification of perivascular fibrosis (**e**) at 8 weeks in male C57Bl/6J sham (*n=8*), TAC controls (*n=5-8*); and anti-VEGFR3-treated TAC (*n=7*). Examples of perivascular lymphatics in the interventricular septum (**g**) at 3 weeks post-TAC, visualized by light sheet microscopy. White dashed arrow denotes localization of the septal artery. αSMA, *red*; Lyve1, *cyan*. (Scale bar, 500 µm.) Groups were compared by non-parametric Kruskal Wallis followed by Dunn’s posthoc test (for histology) and by two-way ANOVA followed by Dunnett’s multiple comparisons test (for expression analyses). * p<0.05, ** p<0.01, *** p<0.001 versus sham; ## p<0.01, ### p<0.001 *versus* control TAC.

Next, we investigated cardiac fibrosis at 3- and 8-weeks post-TAC in male C57Bl/6J mice treated or not with anti-VEGFR3. Using cardiac gene expression analysis, we found that the levels of the pro-fibrotic cytokine *Il-6* and collagen 1, but not collagen 3 (*Col1a1* and *Col3a1* genes), were similarly increased at 8 weeks post-TAC in both anti-VEGFR3-treated and control IgG-treated mice, as compared to sham (**Fig. 6c**). Interstitial cardiac fibrosis, evaluated by either collagen 1 or collagen 3 immunohistochemistry, was also similarly increased in both TAC groups (**Fig. 6f, Suppl. Fig. 8a, b**). In contrast, in line with our findings in female C57Bl/6J mice, the level of perivascular fibrosis was found to be significantly increased in anti-VEGFR3-treated mice at 8 weeks post-TAC (**Fig. 6b, e**). Examining lymphatics in the region of the septal artery, we found that the pressure-overload led to an expansion of perivascular lymphatics, which was prevented by anti-VEGFR3 treatment (**Fig. 6g**). Taken together, this indicates that poor lymphangiogenesis represent as potential risk factor for perivascular, but not interstitial, fibrosis during pressure-overload, potentially linked to insufficient lymphatic drainage of the immediate surroundings of hypertensive arteries.

## Discussion

In this study, we demonstrate that pressure-overload in mice leads to gender- and strain-dependent stimulation of lymphangiogenesis through cardiac upregulation of lymphangiogenic growth factors (**Fig. 7**). Our results are in agreement with previous reports of increased Vegfc and Vegfd expression during cardiac hypertrophy in mice and men ^30–32^. Additionally, our data indirectly suggest that pro-inflammatory cytokines, especially IL-1β, may further increase cardiac *Vegfc* expression to stimulate robust expansion of the lymphatic network during pressure-overload. Conversely, we show that inhibition of lymphangiogenesis post-TAC led to aggravation of cardiac inflammation, hypertrophy and perivascular fibrosis, and accelerated the development of cardiac dysfunction and adverse remodeling.

**Fig. 7.**
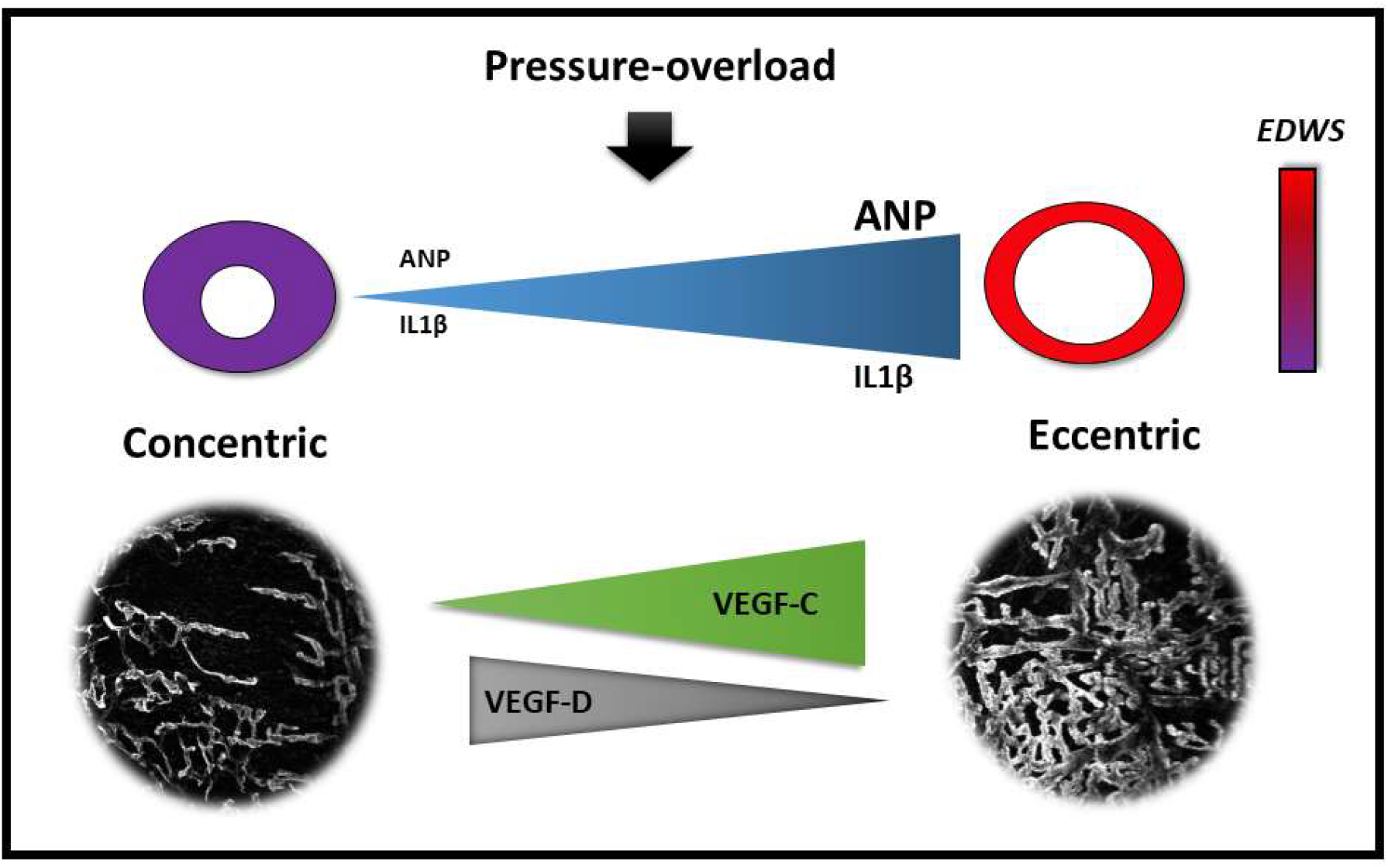
Schematic overview. Pressure-overload leads to strain-dependent differences in the cardiac response in mice, with either concentric hypertrophy or eccentric cardiac hypertrophy accompanied by left ventricular dilation causing increased end-diastolic wall stress (*EDWS*). In our study, Balb/c mice developed a dilated phenotype, associated with a strong increase in ANP and IL1β cardiac gene expression. This led to potent stimulation of cardiac lymphangiogenesis, mainly driven by VEGF-C. In contrast, C57Bl/6J mice developed a less dilated phenotype, associated with weaker cardiac upregulation of ANP and Il1β gene expression. This was associated with a weaker, VEGF-D-induced, cardiac lymphangiogenic response. Nevertheless, inhibition of this lymphangiogenesis during pressure-overload led to aggravation of cardiac inflammation and perivascular fibrosis, accelerating the development of cardiac remodeling and decompensation.

Our indirect estimates, based on evaluation of cardiac *Nppa* and *Nppb* expression levels, indicate severely elevated wall stress (despite similarly increased aortic pressures post-TAC) in Balb/c versus C57BL/6J females, both initially and at the end of the study. We propose that one key to the difference observed in lymphangiogenesis following pressure-overload may be the development of a concentric, rather than eccentric, cardiac hypertrophic phenotype in C57BL/6J females. This limits wall stress, given that stress correlates positively with left ventricular *pressure* (similar increase post-TAC in both strains) and *chamber size* (larger post-TAC in Balb/c vs female C57Bl/6J). Indeed, wall stress, as estimated by cardiac *Nppa* expression, was considerably lower by 6-8 weeks post-TAC in female C57Bl/6J as compared to Balb/c. We postulate that this may have contributed to reduce the lymphangiogenic drive after the initially increased cardiac Vegfd expression seen in C57Bl/6J females. In contrast, in Balb/c mice, the wall stress increased throughout the study, as left ventricular dilation (44% increase in LV ESD at 8 weeks post-TAC *vs* sham) was accompanied by ventricular wall thinning (15-20% reduction in AWT ES and PWT ES, **Suppl. table 2**). It thus seems that increased EDWS (due to ventricular dilation), rather than wall thickening (due to cardiac hypertrophy), may be the main trigger for induction of cardiac lymphangiogenesis during pressure-overload. Interestingly, this may hold true also in humans, as we observed increased lymphatic density in patients with DCM as compared to healthy controls. In parallel, a clinical study in HF patients indicated that arterial wall stress, due to elevated pulmonary artery wedge pressure, increased plasma Vegfd levels ^33^.

The penetration depth into the myocardium of cardiac lymphatics is species-dependent, with more intramyocardial branches observed in larger animals with thicker walls. For example, previous studies in human and dog hearts ^34,35^ indicate that the lymphatic network penetrates the entire myocardium in these species. In contrast, mice, with left ventricular wall thickness of around 1 mm, display lymphatic vessels essentially limited to the cardiac surface also following pressure-overload (**suppl. video**). We speculate that this differential species-dependent lymphatic penetration, likely influenced by the level of ventricular wall stress (rather than wall thickness *per se*) may reflect increased metabolic needs to clear waste products from the cardiac interstitium under elevated EDWS settings. An additional component that raises EDWS is myocardial edema, which increases the diastolic stress-strain relationship and decreases cardiac compliance ^36^. However, comparing Balb/c and C57Bl/6J females, we found that the myocardial edema was similar post-TAC, while our indirect estimates of EDWS indicated much higher stress in the thinning walls of Balb/c mice.

We previously demonstrated that therapeutic lymphangiogenesis limited myocardial edema post-MI in rats ^6^. It should be noted that following MI there is significant destruction, rarefaction, and dysfunction of cardiac lymphatics, whereas after pressure-overload cardiac lymphatics were comparably less altered. In particular, we recently demonstrated that poor cardiac lymphangiogenesis post-MI was linked to elevated interferon (IFN)-γ, in part produced by cardiac-infiltrating T cells ^8^. In contrast, previous studies have indicated a predominant Th2-type immune response in Balb/c mice, with low cardiac expression of IFN-γ and elevated IL-4, leading to dilated cardiomyopathy in response to chronic arterial hypertension ^37^. Potentially, this immune phenotype of low IFN-γ and elevated IL-1β in the heart of Balb/c mice may have been conductive to the robust lymphangiogenic response observed following pressure-overload. In this context, we expected more rapid restoration of myocardial fluid balance post-TAC in Balb/c versus in C57Bl/6J mice. Thus, our findings that myocardial edema was similarly severe in all three TAC models, characterized by different degrees of cardiac lymphangiogenesis, was surprising. However, the expansion of essentially capillary lymphatics in Balb/c was clearly not sufficient to restore fluid balance. Alternatively, it is possible that these newly formed lymphatics were non-functional and leaky, and thus inept to restore fluid homeostasis. Moreover, given that we only observed a tendency for an increase in myocardial water content in the anti-VEGFR3 treated TAC group that displayed significantly reduced lymphatic density, we postulate that the main culprit of cardiac edema in our model may be unrelated to lymphangiogenesis, and instead reflect blood vascular hyperpermeability in the heart resulting from increased coronary blood pressure after aortic banding. Thus, the increased influx of plasma-derived fluids effectively overwhelmed the drainage capacity of cardiac lymphatics, even after potent lymphangiogenesis as in Balb/c. Importantly, this setting is specific to the TAC model, and does not apply in patients with aortic stenosis, who suffer from reduced, rather than increased, perfusion of the coronary vasculature. Thus, it remains to be determined, in a more physiologically-relevant model, whether robust cardiac lymphangiogenesis in response to pressure-overload may be sufficient to improve myocardial fluid balance.

Our data promisingly revealed that moderate cardiac lymphangiogenesis, including periarterial lymphatic expansion, sufficed to limit perivascular fibrosis post-TAC in male C57Bl/6J. However, we were surprised by the absence of impact of anti-VEGFR3 treatment on interstitial fibrosis in C57Bl/6J males, and by the similar level of interstitial fibrosis post-TAC observed in C57Bl/6 vs Balb/c females. This differs from our previous findings post-MI in rats, where lymphangiogenesis reduced the development of interstitial fibrosis ^6^. However, one important difference is the degree of cardiac hypoxia and inflammation, both potent drivers of fibrosis ^38^, which both are more severe post-MI as compared to following pressure-overload. However, given that ventricular wall stress, as well as cardiac expression of pro-fibrotic *Il1β* and *Il6* cytokines, was higher in Balb/c versus female C57Bl/6J, it seems impertinent to attempt any direct conclusion on how lymphangiogenesis may or may not have altered interstitial cardiac fibrosis between these two strains. Indeed, the metabolic milieu of cardiomyocytes and cardiac fibroblasts was likely very different post-TAC in these two strains due to the difference in EDWS ^39^.

During the preparation of this manuscript, another study reported on the beneficial effects of therapeutic lymphangiogenesis in a mouse pressure-overload model ^40^. The authors surprisingly demonstrate that systemic daily injections of recombinant Vegfc, an inefficient therapeutic modality ^8,10^, potently stimulated cardiac lymphangiogenesis and almost completely prevented myocardial edema, cardiac hypertrophy and dysfunction in their model. It should be noted that their study was based on a single-banding TAC method, shown to produce non-permanent constriction in up to 30% of animals ^16^. Although the authors argue that the lower cardiac *Nppa* expression observed after their Vegfc therapy was a sign of the cardiac benefit, it is challenging to understand how their systemic therapeutic approach so fully protected against the deleterious effects of pressure-overload as to both prevent and revert ventricular dilation and hypertrophy. Our study reveals a subtler functional impact of cardiac lymphangiogenesis during pressure-overload, even in the setting of extensive expansion of the lymphatic network, as observed in female Balb/c. Indeed, although we speculate that the endogenous lymphangiogenesis may have limited cardiac inflammation and perivascular fibrosis in Balb/c, these mice were not protected against either chronic myocardial edema or cardiac decompensation. On the other hand, the moderate lymphangiogenic response seen in C57Bl/6J male mice was sufficient to limit cardiac inflammation and perivascular fibrosis, and to slow the development of cardiac hypertrophy and decompensation. We hypothesize that this functional cardiac difference between C57Bl/6J and Balb/c mice in response to the same pressure-overload may reflect differing immune mechanisms underlying ventricular dilation in these two strains, with more severe dilation linked to the elevated Il1β in Balb/c mice.

In conclusion, our findings of elevated lymphatic density in mice with dilated cardiac phenotype, as well as in HF patients with DCM, indicate that ventricular dilation, rather than cardiac hypertrophy, triggers lymphatic expansion in the heart. Future studies will reveal whether EDWS is directly linked to lymphangiogenic growth factor expression in the human heart, as suggested by our experimental study. As for the clinical outlook, it remains to be determined whether endogenous differences in cardiac lymphangiogenic potential in patients with aortic stenosis or chronic arterial hypertension may impact cardiac remodeling and HF development. Our work suggests that individuals with poor cardiac expression of lymphatic growth factors, either Vegfc or Vegfd, may be at increased risk for aggravation of cardiac inflammation and perivascular fibrosis, leading to accelerated progression of cardiac dysfunction during pressure-overload.

## Author contributions

C.H, A.D, M.H, H.C and O.H performed and analyzed immunohistochemistry and histology. C.H and A.D performed and analyzed flow cytometry with guidance of G.R and V.T. Echocardiography was performed and analyzed by C.H, M.H, M.D and O.H; microgravimetry was carried out by C.H, O.H and E.B; A.L, D.G, D.S. and R.H carried-out, post-processed and analyzed light sheet and confocal imaging; JP.H. carried out surgical mouse model; C.H, S.R and M.H carried-out cardiac gene expression analyses with guidance of S.F; JB.M managed the biobank at Hopital Bichat and contributed clinical data for this study; V.T, P.M, A.O.P and V.R participated in manuscript preparation; E.B and C.H. designed the study, analyzed results, and prepared the manuscript draft. All authors approved of the final version of the manuscript.

## Acknowledgments

We thank Manon Douyère, Frederic Rabin, and Théo Lemarcis at Normandy University, France for assistance with immunohistochemistry and RTqPCR, and Pr. Anna Ratajska at the Department of Pathology, Medical University of Warsaw, Center for Biostructure, Poland, for helpful discussions.

## References

1. Brakenhielm E, Alitalo K. Cardiac lymphatics in health and disease. Nat Rev Cardiol. 2019;16:56–68.

2. Jalkanen S, Salmi M. Lymphatic endothelial cells of the lymph node. Nat Rev Immunol. 2020;20:566–578.

3. Nahrendorf M. Myeloid cell contributions to cardiovascular health and disease. Nat Med. 2018;24:711–720.

4. Van der Borght K, Scott CL, Nindl V, Bouché A, Martens L, Sichien D, Van Moorleghem J, Vanheerswynghels M, De Prijck S, Saeys Y, Ludewig B, Gillebert T, Guilliams M, Carmeliet P, Lambrecht BN. Myocardial Infarction Primes Autoreactive T Cells through Activation of Dendritic Cells. Cell Rep. 2017;18:3005–3017.

5. Sintou A, Mansfield C, Iacob A, Chowdhury RA, Narodden S, Rothery SM, Podovei R, Sanchez-Alonso JL, Ferraro E, Swiatlowska P, Harding SE, Prasad S, Rosenthal N, Gorelik J, Sattler S. Mediastinal Lymphadenopathy, Class-Switched Auto-Antibodies and Myocardial Immune-Complexes During Heart Failure in Rodents and Humans. Front Cell Dev Biol. 2020;8:695.

6. Henri O, Pouehe C, Houssari M, Galas L, Nicol L, Edwards-Lévy F, Henry J-P, Dumesnil A, Boukhalfa I, Banquet S, Schapman D, Thuillez C, Richard V, Mulder P, Brakenhielm E. Selective Stimulation of Cardiac Lymphangiogenesis Reduces Myocardial Edema and Fibrosis Leading to Improved Cardiac Function Following Myocardial Infarction. Circulation. 2016;133:1484–1497.

7. Vieira JM, Norman S, Villa Del Campo C, Cahill TJ, Barnette DN, Gunadasa-Rohling M, Johnson LA, Greaves DR, Carr CA, Jackson DG, Riley PR. The cardiac lymphatic system stimulates resolution of inflammation following myocardial infarction. J Clin Invest. 2018;128:3402–3412.

8. Houssari M, Dumesnil A, Tardif V, Kivelä R, Pizzinat N, Boukhalfa I, Godefroy D, Schapman D, Hemanthakumar KA, Bizou M, Henry J-P, Renet S, Riou G, Rondeaux J, Anouar Y, Adriouch S, Fraineau S, Alitalo K, Richard V, Mulder P, Brakenhielm E. Lymphatic and Immune Cell Cross-Talk Regulates Cardiac Recovery After Experimental Myocardial Infarction. Arterioscler Thromb Vasc Biol. 2020;40:1722–1737.

9. Yang G-H, Zhou X, Ji W-J, Zeng S, Dong Y, Tian L, Bi Y, Guo Z-Z, Gao F, Chen H, Jiang T-M, Li Y-M. Overexpression of VEGF-C attenuates chronic high salt intake-induced left ventricular maladaptive remodeling in spontaneously hypertensive rats. Am J Physiol Heart Circ Physiol. 2014;306:H598–609.

10. Song L, Chen X, Swanson TA, LaViolette B, Pang J, Cunio T, Nagle MW, Asano S, Hales K, Shipstone A, Sobon H, Al-Harthy SD, Ahn Y, Kreuser S, Robertson A, Ritenour C, Voigt F, Boucher M, Sun F, Sessa WC, Roth Flach RJ. Lymphangiogenic therapy prevents cardiac dysfunction by ameliorating inflammation and hypertension. eLife. 2020;9.

11. Beaini S, Saliba Y, Hajal J, Smayra V, Bakhos J-J, Joubran N, Chelala D, Fares N. VEGF-C attenuates renal damage in salt-sensitive hypertension. J Cell Physiol. 2019;234:9616–9630.

12. Tomanek RJ. Response of the coronary vasculature to myocardial hypertrophy. J Am Coll Cardiol. 1990;15:528–533.

13. Zheng W, Seftor EA, Meininger CJ, Hendrix MJ, Tomanek RJ. Mechanisms of coronary angiogenesis in response to stretch: role of VEGF and TGF-beta. Am J Physiol Heart Circ Physiol. 2001;280:H909–917.

14. Sano M, Minamino T, Toko H, Miyauchi H, Orimo M, Qin Y, Akazawa H, Tateno K, Kayama Y, Harada M, Shimizu I, Asahara T, Hamada H, Tomita S, Molkentin JD, Zou Y, Komuro I. p53-induced inhibition of Hif-1 causes cardiac dysfunction during pressure overload. Nature. 2007;446:444–448.

15. Hu P, Zhang D, Swenson L, Chakrabarti G, Abel ED, Litwin SE. Minimally invasive aortic banding in mice: effects of altered cardiomyocyte insulin signaling during pressure overload. Am J Physiol - Heart Circ Physiol. 2003;285:H1261–H1269.

16. Lygate CA, Schneider JE, Hulbert K, Hove M, Sebag-Montefiore LM, Cassidy PJ, Clarke K, Neubauer S. Serial high resolution 3D-MRI after aortic banding in mice: band internalization is a source of variability in the hypertrophic response. Basic Res Cardiol. 2006;101:8–16.

17. Burton JB, Priceman SJ, Sung JL, Brakenhielm E, An DS, Pytowski B, Alitalo K, Wu L. Suppression of prostate cancer nodal and systemic metastasis by blockade of the lymphangiogenic axis. Cancer Res. 2008;68:7828–7837.

18. Yamakawa H, Imamura T, Matsuo T, Onitsuka H, Tsumori Y, Kato J, Kitamura K, Koiwaya Y, Eto T. Diastolic wall stress and ANG II in cardiac hypertrophy and gene expression induced by volume overload. Am J Physiol Heart Circ Physiol. 2000;279:H2939–2946.

19. Vanderheyden M, Goethals M, Verstreken S, De Bruyne B, Muller K, Van Schuerbeeck E, Bartunek J. Wall stress modulates brain natriuretic peptide production in pressure overload cardiomyopathy. J Am Coll Cardiol. 2004;44:2349–2354.

20. Ristimäki A, Narko K, Enholm B, Joukov V, Alitalo K. Proinflammatory cytokines regulate expression of the lymphatic endothelial mitogen vascular endothelial growth factor-C. J Biol Chem. 1998;273:8413–8418.

21. Weichand B, Popp R, Dziumbla S, Mora J, Strack E, Elwakeel E, Frank A-C, Scholich K, Pierre S, Syed SN, Olesch C, Ringleb J, Ören B, Döring C, Savai R, Jung M, von Knethen A, Levkau B, Fleming I, Weigert A, Brüne B. S1PR1 on tumor-associated macrophages promotes lymphangiogenesis and metastasis via NLRP3/IL-1β. J Exp Med. 2017;214:2695–2713.

22. Bletsa A, Abdalla H, Løes S, Berggreen E. Lymphatic growth factors are expressed in human gingiva and upregulated in gingival fibroblasts after stimulation. J Periodontol. 2018;89:606–615.

23. Morfoisse F, Tatin F, Chaput B, Therville N, Vaysse C, Métivier R, Malloizel-Delaunay J, Pujol F, Godet A-C, De Toni F, Boudou F, Grenier K, Dubuc D, Lacazette E, Prats A-C, Guillermet-Guibert J, Lenfant F, Garmy-Susini B. Lymphatic Vasculature Requires Estrogen Receptor-α Signaling to Protect From Lymphedema. Arterioscler Thromb Vasc Biol. 2018;38:1346–1357.

24. Damilano F, Franco I, Perrino C, Schaefer K, Azzolino O, Carnevale D, Cifelli G, Carullo P, Ragona R, Ghigo A, Perino A, Lembo G, Hirsch E. Distinct effects of leukocyte and cardiac phosphoinositide 3-kinase γ activity in pressure overload-induced cardiac failure. Circulation. 2011;123:391–399.

25. Laroumanie F, Douin-Echinard V, Pozzo J, Lairez O, Tortosa F, Vinel C, Delage C, Calise D, Dutaur M, Parini A, Pizzinat N. CD4+ T cells promote the transition from hypertrophy to heart failure during chronic pressure overload. Circulation. 2014;129:2111–2124.

26. Kallikourdis M, Martini E, Carullo P, Sardi C, Roselli G, Greco CM, Vignali D, Riva F, Ormbostad Berre AM, Stølen TO, Fumero A, Faggian G, Di Pasquale E, Elia L, Rumio C, Catalucci D, Papait R, Condorelli G. T cell costimulation blockade blunts pressure overload-induced heart failure. Nat Commun. 2017;8:14680.

27. Ma X-L, Lin Q-Y, Wang L, Xie X, Zhang Y-L, Li H-H. Rituximab prevents and reverses cardiac remodeling by depressing B cell function in mice. Biomed Pharmacother Biomedecine Pharmacother. 2019;114:108804.

28. Martini E, Kunderfranco P, Peano C, Carullo P, Cremonesi M, Schorn T, Carriero R, Termanini A, Colombo FS, Jachetti E, Panico C, Faggian G, Fumero A, Torracca L, Molgora M, Cibella J, Pagiatakis C, Brummelman J, Alvisi G, Mazza EMC, Colombo MP, Lugli E, Condorelli G, Kallikourdis M. Single-Cell Sequencing of Mouse Heart Immune Infiltrate in Pressure Overload-Driven Heart Failure Reveals Extent of Immune Activation. Circulation. 2019;140:2089–2107.

29. Brakenhielm E, González A, Díez J. Role of Cardiac Lymphatics in Myocardial Edema and Fibrosis: JACC Review Topic of the Week. J Am Coll Cardiol. 2020;76:735–744.

30. Abraham D, Hofbauer R, Schäfer R, Blumer R, Paulus P, Miksovsky A, Traxler H, Kocher A, Aharinejad S. Selective downregulation of VEGF-A(165), VEGF-R(1), and decreased capillary density in patients with dilative but not ischemic cardiomyopathy. Circ Res. 2000;87:644–647.

31. Kholová I, Dragneva G, Cermáková P, Laidinen S, Kaskenpää N, Hazes T, Cermáková E, Steiner I, Ylä-Herttuala S. Lymphatic vasculature is increased in heart valves, ischaemic and inflamed hearts and in cholesterol-rich and calcified atherosclerotic lesions. Eur J Clin Invest. 2011;41:487–497.

32. Huusko J, Lottonen L, Merentie M, Gurzeler E, Anisimov A, Miyanohara A, Alitalo K, Tavi P, Ylä-Herttuala S. AAV9-mediated VEGF-B gene transfer improves systolic function in progressive left ventricular hypertrophy. Mol Ther J Am Soc Gene Ther. 2012;20:2212–2221.

33. Houston BA, Tedford RJ, Baxley RL, Sykes B, Powers ER, Nielsen CD, Steinberg DH, Maran A, Fernandes VLC, Todoran T, Jones JA, Zile MR. Relation of Lymphangiogenic Factor Vascular Endothelial Growth Factor-D to Elevated Pulmonary Artery Wedge Pressure. Am J Cardiol. 2019;124:756–762.

34. Shore LR. The lymphatic drainage of the human heart. J Anat. 1929;63:291.

35. Patek, P. The morphology of the lymphatics of the mammalian heart. Am. J Anat. 1939;203–249.

36. Dongaonkar RM, Stewart RH, Quick CM, Uray KL, Cox CS, Laine GA. Award article: Microcirculatory Society Award for Excellence in Lymphatic Research: time course of myocardial interstitial edema resolution and associated left ventricular dysfunction. Microcirc N Y N 1994. 2012;19:714–722.

37. Peng H, Yang X-P, Carretero OA, Nakagawa P, D’Ambrosio M, Leung P, Xu J, Peterson EL, González GE, Harding P, Rhaleb N-E. Angiotensin II-induced dilated cardiomyopathy in Balb/c but not C57BL/6J mice. Exp Physiol. 2011;96:756–764.

38. López B, Ravassa S, Moreno MU, José GS, Beaumont J, González A, Díez J. Diffuse myocardial fibrosis: mechanisms, diagnosis and therapeutic approaches. Nat Rev Cardiol. 2021;

39. Kissling G. Mechanical determinants of myocardial oxygen consumption with special reference to external work and efficiency. Cardiovasc Res. 1992;26:886–892.

40. Lin Q, Zhang Y, Bai J, Liu J, Li H. VEGF-C/VEGFR-3 axis protects against pressure-overload induced cardiac dysfunction through regulation of lymphangiogenesis. Clin Transl Med [Internet]. 2021 [cited 2021 Mar 30];11. Available from: https://onlinelibrary.wiley.com/doi/10.1002/ctm2.374

