## Supplementary material for "Regulation and impact of cardiac lymphangiogenesis in pressure-overload-induced heart failure": Suppl. figures and methods

### Supplementary Figures

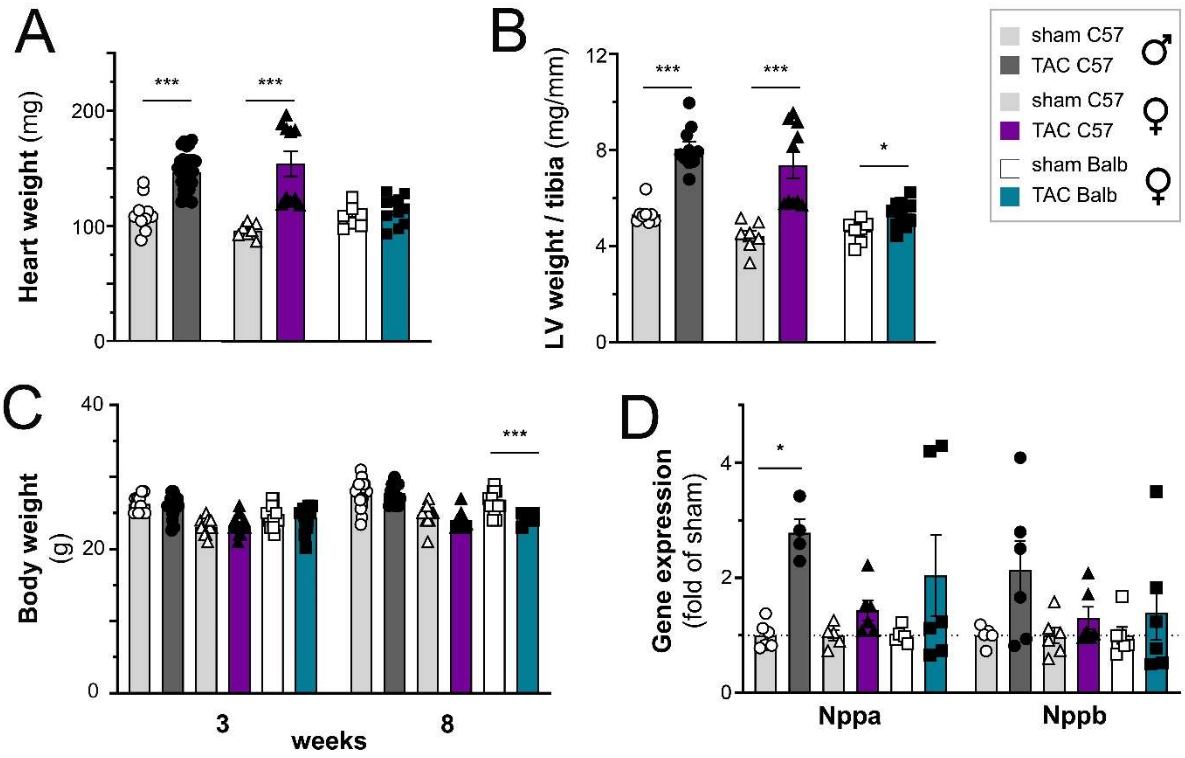


S1 **Evaluation of cardiac hypertrophy at 3 weeks post-TAC**

Morphometric assessment of heart weight (**a**), left ventricular (LV) weight normalized to tibia lengths (**b**) at 3 weeks in C57Bl/6J healthy male sham (open circles, *n=7-11*), post-TAC males (closed circles, *n=10-29*), healthy female sham (open triangles, *n*=7), post-TAC females (closed triangles, *n=10*), and in Balb/c females either sham (open square, *n*=7) or post-TAC (closed square, *n=11*). Assessment of body weight gain throughout the study (**c**). Cardiac expression analyses of *Nppa* and *Nppb* (*n=6* per group) at 3 weeks (**d**). Groups were compared pair-wise by two-way ANOVA followed by Sidak's multiple comparisons. * $p<0.05$, *** $p<0.001$.

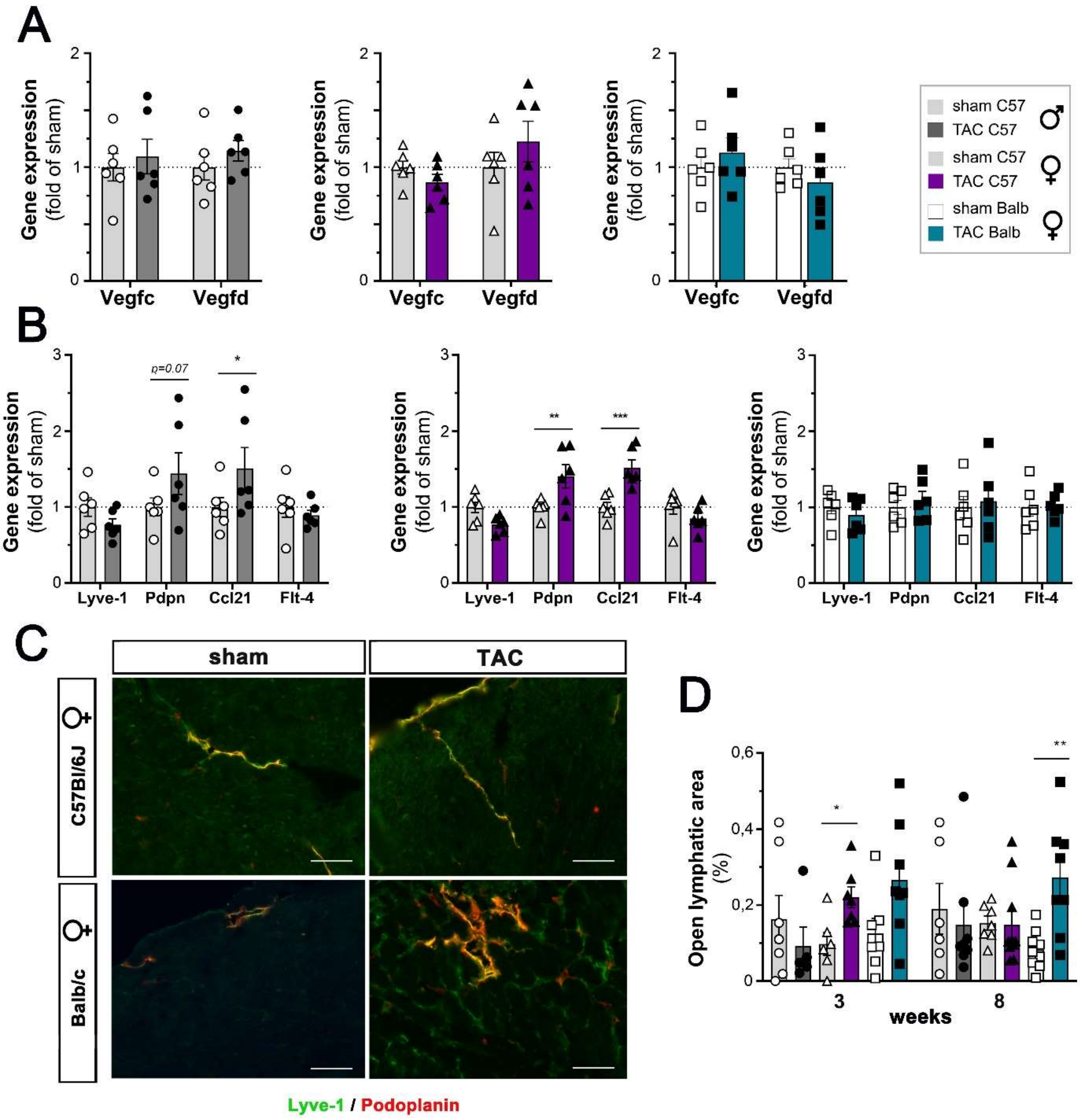


#### S2 Evaluation of cardiac lymphatics at 3 and 8 weeks post-TAC

Cardiac expression analyses of Vegfc and Vegfd (**a**) and lymphatic markers *Lyve1, Pdpn, Ccl21*, and *Flt4* (**b**) at 3 weeks in C57Bl/6J healthy male sham (open circles, *n=6*), post-TAC males (closed circles, *n=6*), healthy female sham (open triangles, *n=6*), post-TAC females (closed triangles, *n=6*), and in Balb/c females either sham (open square, *n=6*) or post-TAC (closed square, *n=6*). Examples (**c**) of immunohistochemical analysis at 8 weeks of lymphatic markers Lyve1 (*green*) and Podoplanin (*red*) in cardiac sections imaged at x20. (Scale bar 50 µm). Quantification of open lymphatic area per heart (**d**) at 3 and 8 weeks post-TAC (*n=5-10* animals per group). Groups were compared by one-way ANOVA followed by Bonferroni posthoc test (for immunohistochemical analyses) and by two-way ANOVA followed by Dunnett's multiple comparisons test (for expression analyses). * p<0.05, ** p<0.01, *** p<0.001.

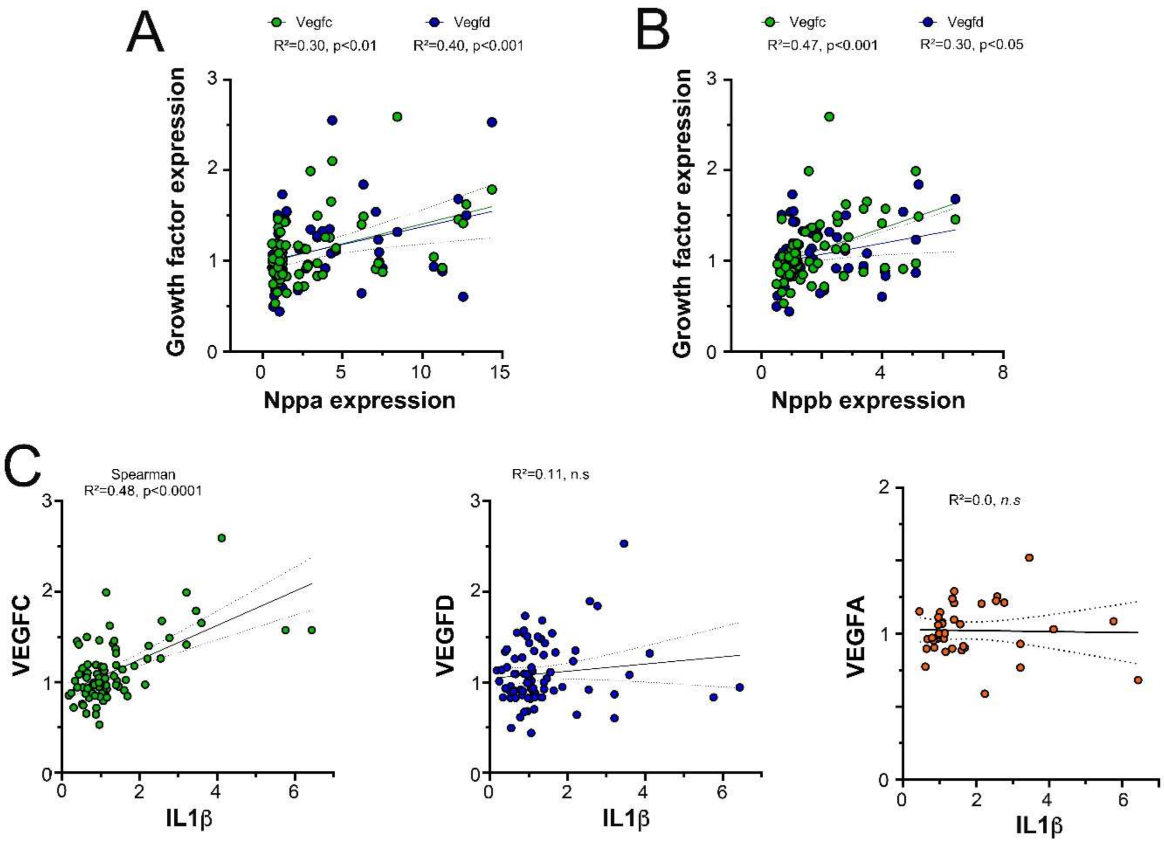


S3 **Correlation analysis for lymphangiogenic growth factors**

Cardiac gene expression of *Vegfc* (*green circles*) and *Vegfd* (*blue circles*) correlated with both cardiac *Nppa* (**a**) and *Nppb* (**b**). In contrast, only *Vegfc* correlated with cardiac *IL1β* gene expression (**c**). Non-parametric Spearman correlation analysis.

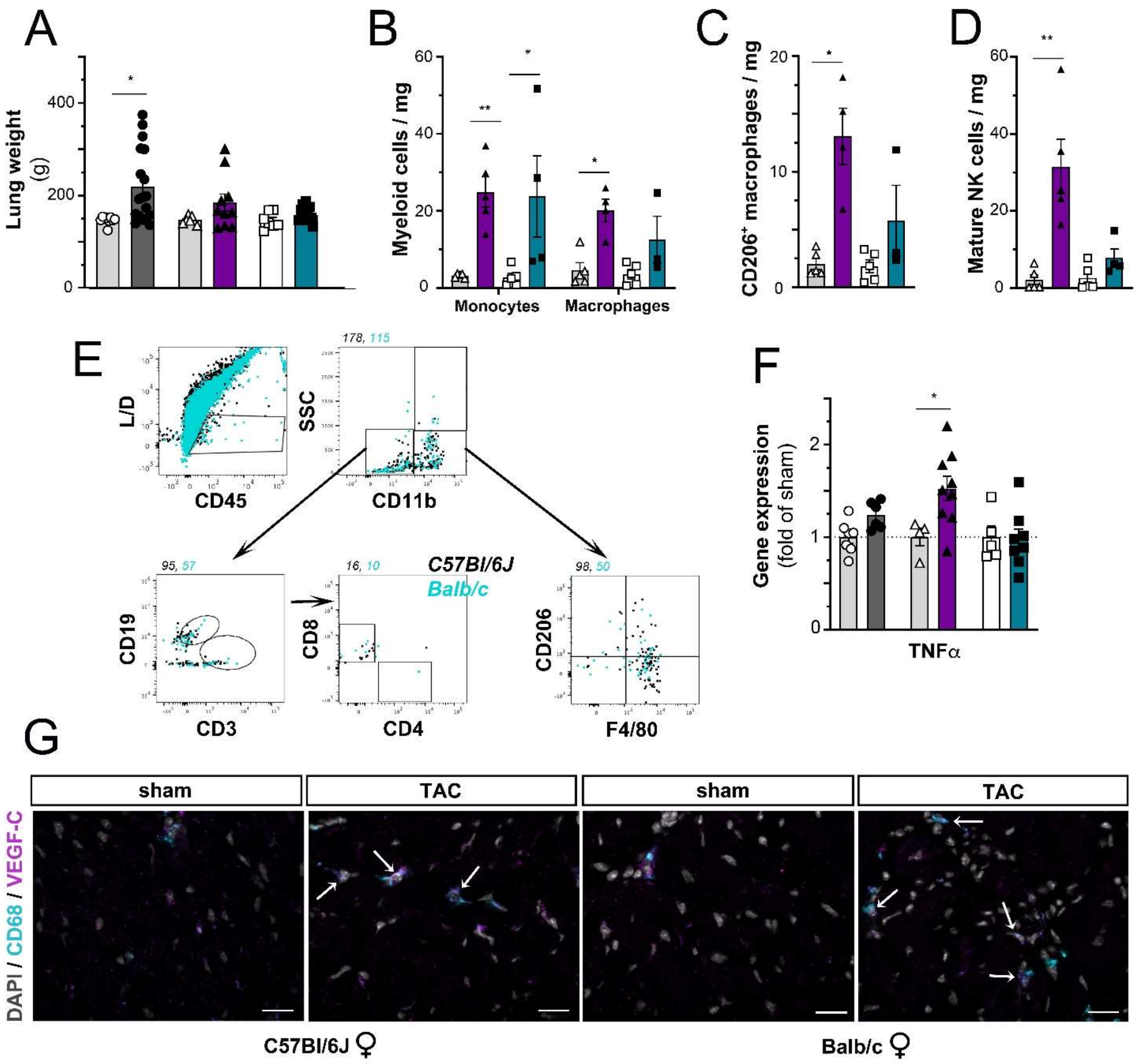


##### S4 Assessment of cardiac inflammation and pulmonary edema post-TAC

Evaluation of pulmonary weights at 3 weeks (**a**) in C57Bl/6J males: sham (open circles, *n=8*) or TAC (closed circles, *dark grey bar n=16*); C57Bl/6J females: sham (open triangles, *n=5*) or TAC (closed triangles, *purple bar n=10*); and in Balb/c females: sham (open square, *n=5*) or TAC (closed square, *blue bar n=10*). Evaluation at 8 weeks post-TAC of cardiac-infiltrating immune cells (*n=4-5 samples* per group) including CD11b$^{+}$ SSC$^{low}$ monocytes and F4-80$^{+}$ macrophages (**b**), CD206$^{+}$ alternative macrophage subpopulation (**c**), and mature NK1.1$^{+}$ CD19$^{-}$ CD11b$^{+}$ NK cells (**d**). Examples of flow cytometry gating in healthy sham mice (**e**). Numbers indicate total events per group in the gate. Cardiac gene expression analysis of *Tnfα* (**f**) at 8 weeks post-TAC (*n=5-10* animals per group). Examples of Vegfc-expressing cardiac macrophages, indicated by arrows, at 3 weeks post-TAC visualized by immunohistochemistry (Vegfc, *magenta*; CD68, *cyan*; DAPI, grey *x40, scalebar 20* µm) (**g**). Groups were compared pair-wise by non-parametric Kruskal Wallis followed by Dunn's posthoc test (gene expression analysis). * $p<0.05$, ** $p<0.01$.

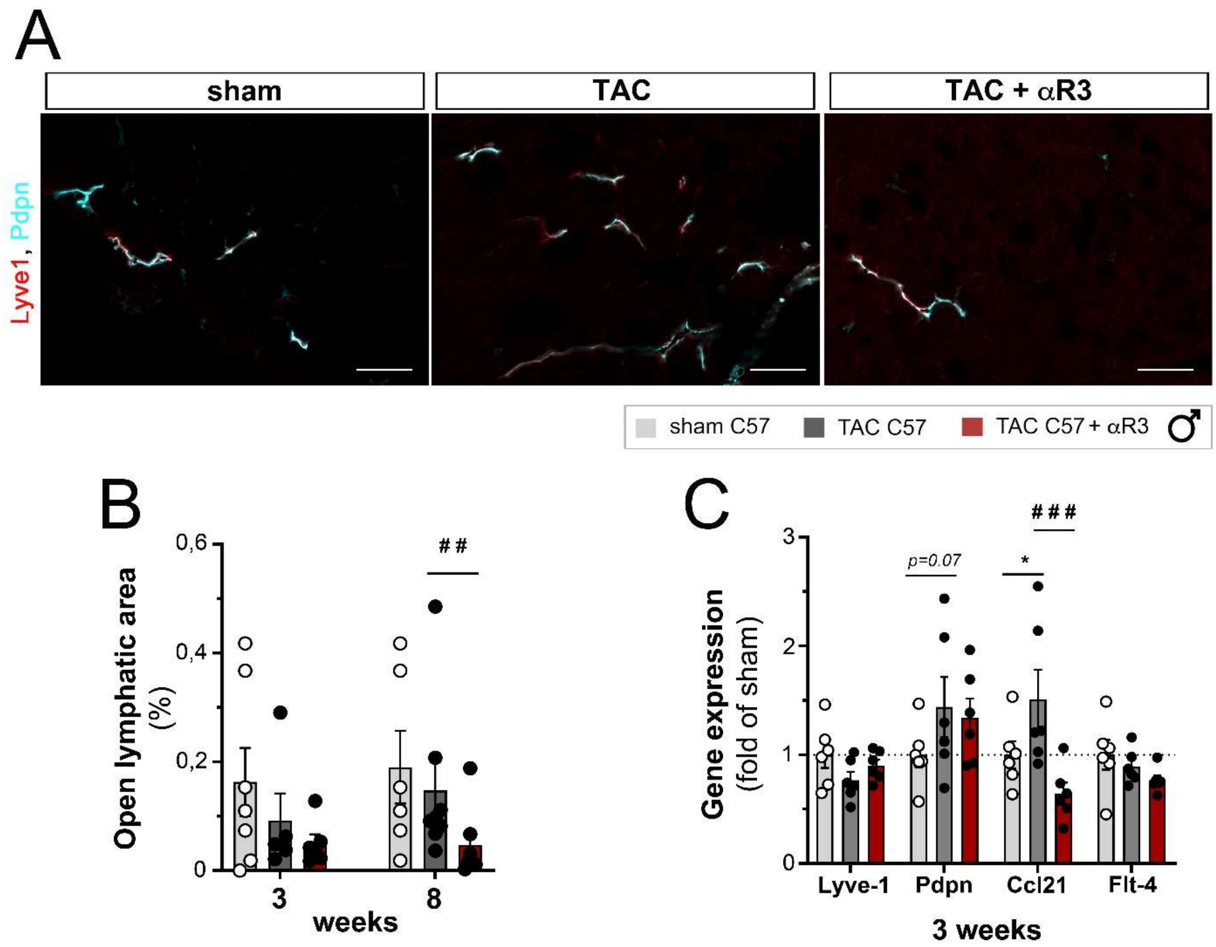


S5 **Anti-VEGFR3 treatment blocks cardiac lymphangiogenesis post-TAC**

Examples (**a**) of immunohistochemical analysis at 8 weeks post-TAC of lymphatic markers Lyve1 (*red*) and Podoplanin (*blue*) in cardiac sections imaged at *x20*. (Scale bar 20 µm). Quantification of open lymphatic area per heart (**b**) at 3 or 8 weeks post-TAC (*n=5-10* animals per group). Cardiac expression analyses of lymphatic markers *Lyve1, Pdpn, Ccl21*, and *Flt4* (**c**) at 3 weeks in male C57Bl/6J sham mice (open circles, *n=6*), TAC controls (closed circles, *dark grey bar n=6*), and TAC treated with anti-VEGFR3 (closed circles, *red bar n=6*). Groups were compared by non-parametric Kruskal Wallis followed by Dunn's posthoc test (for immunohistochemistry) and by two-way ANOVA followed by Dunnett's multiple comparisons test (for expression analyses). * $p<0.05$ *versus* sham, ## $p<0.01$, ### $p<0.001$ *versus* control TAC.

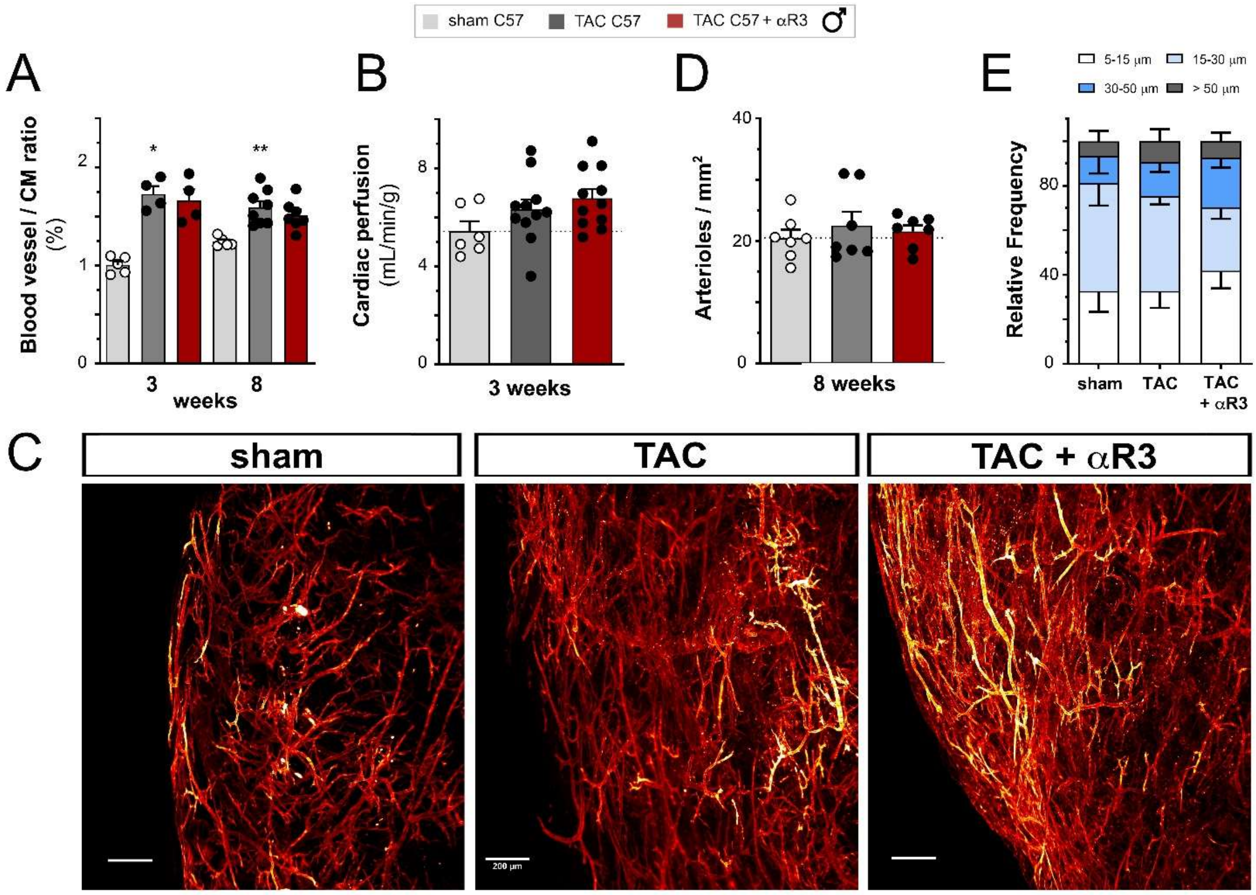


S6 **No effect on angiogenesis or arteriogenesis by anti-VEGFR3 treatment**

Quantification of blood vessel to cardiomyocyte ratios (**a**) at 3 or 8 weeks post-TAC (*n=4-10* animals per group). Evaluation of cardiac perfusion by MRI (**b**) at 3 weeks post-TAC (n=6-11 animals per group). Examples of arterial vasculature (**c**) assessed by light sheet imaging of α-SMA-stained hearts at 8 weeks post-TAC. (Scale bar 200 µm). Quantification of cardiac arterial density (**d**) and frequency of arterial size populations (**e**) at 8 weeks in male C57Bl/6J sham mice (open circles, *n=7*), TAC controls (dark grey bar, closed circles, *n=7*), and TAC treated with anti-VEGFR3 (red bar, closed circles, *n=7*). Groups were compared by non-parametric Kruskal Wallis followed by Dunn's posthoc test (immunohistochemistry). * p<0.05, ** p<0.01 *versus* sham.

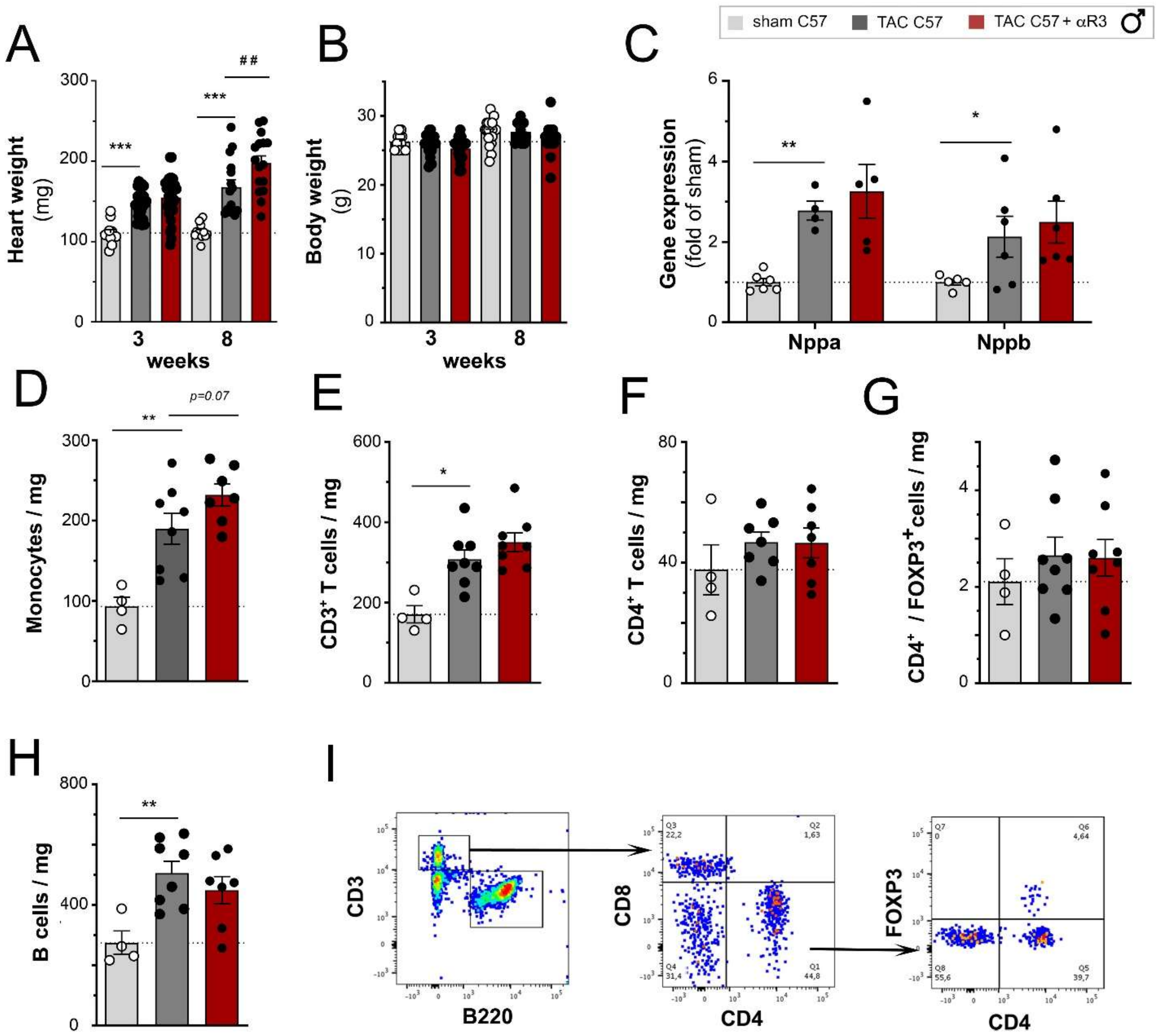


S7 **Anti-VEGFR3 does not increases cardiac B or T cell levels at 3 weeks post-TAC**

Heart weight at 8 weeks (**a**), and body weight gain at 3 and 8 weeks (**b**, *n=11-25* animals per group) in male C57Bl/6J sham mice (open circles), TAC controls (dark grey bar, closed circles), and TAC treated with anti-VEGFR3 (red bar, closed circles). Cardiac gene expression at 3 weeks of *Nppa* and *Nppb* (**c**). Flow cytometric evaluation at 3 weeks post-TAC of cardiac-infiltrating immune cells: CD11b$^{+}$ SSC$^{low}$ monocytes (**d**), CD3$^{+}$ T cells (**e**), CD4$^{+}$ T cells (**f**), CD4$^{+}$/FOXP3$^{+}$ T regulatory cells (**g**), and B220$^{+}$ B cells (**h**). Examples of flow cytometric lymphocyte gating (**i**). Groups were compared by non-parametric Kruskal Wallis followed by Dunn's posthoc test (for flow cytometry) and by two-way ANOVA followed by Dunnett's multiple comparisons test (for body weight and cardiac expression analyses). * p<0.05, ** p<0.01, *** p<0.001 *versus* sham, # p<0.05 *versus* control TAC.

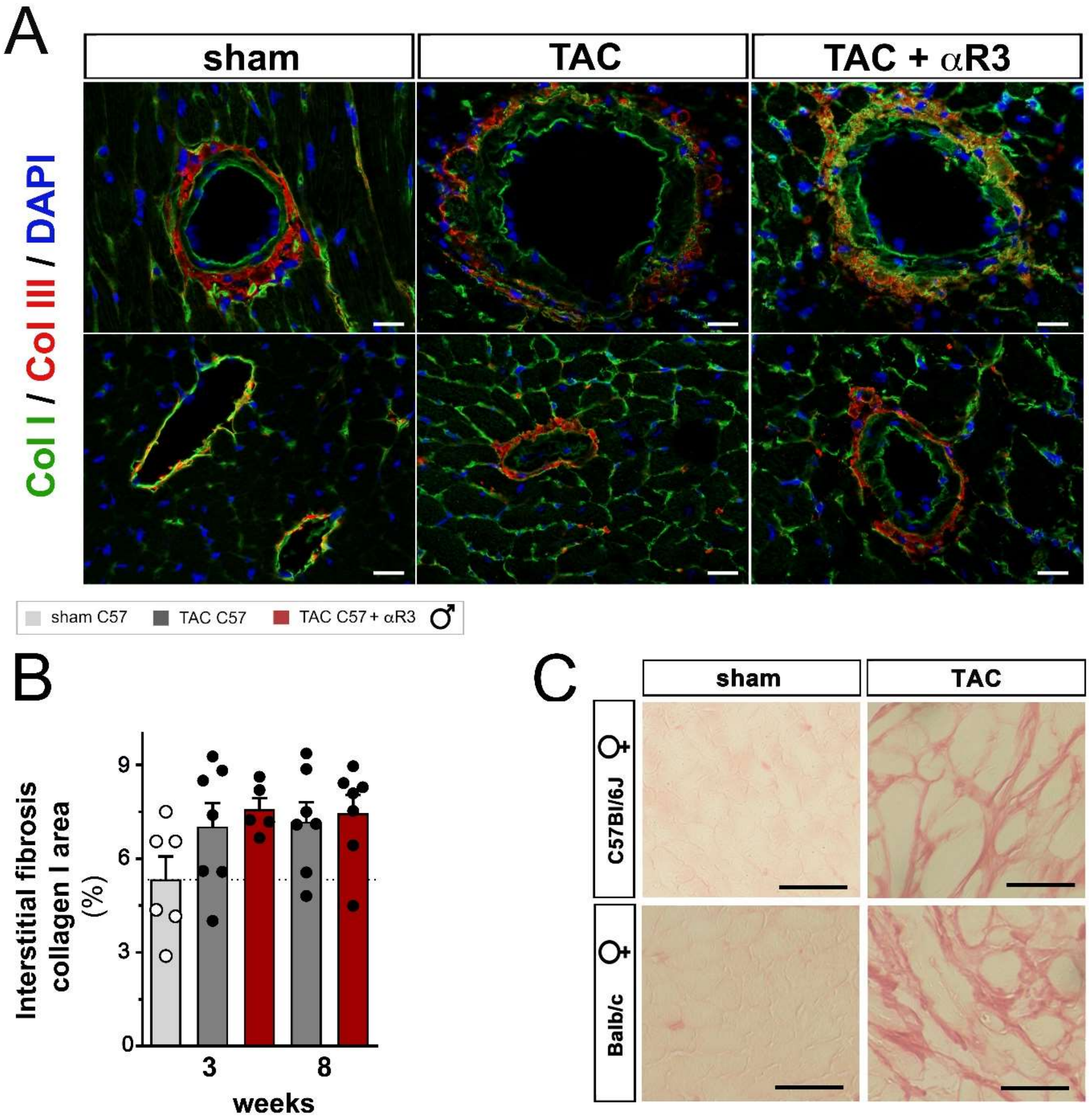


S8 **Cardiac fibrosis evaluated by immunohistochemistry**

Examples of collagen I (*green*) and collagen III (*red*) staining of cardiac sections at 8 weeks (**a**), and quantification of collagen I positive area (**b**, % of total area, *n=5-7* animals per group) in male C57Bl/6J sham mice (open circles), TAC controls (closed circles, *dark grey bar*), and TAC treated with anti-VEGFR3 (closed circles, *red bar*) at 3- or 8-weeks post-TAC. Examples of interstitial fibrosis (**c**) visualized by Sirius red staining of cardiac sections at 8 weeks in female C57Bl/6J and Balb/c mice. (*a*, *x20*, c, *x40*, scale bar 50 µm).

Table S1. **Echocardiography male C57Bl/6J TAC vs sham**

| parameter | | 3 weeks sham | 3 weeks TAC | 3 weeks *vs sham* | 6 weeks sham | 6 weeks TAC | 6 weeks *vs sham* |
|---|---|---|---|---|---|---|---|
| *n* | | *9* | *28* | | *7* | *13* | |
| **BW** | **g** | **26** ± 1 | **25** ± 0 | *n.s* | **27** ± 1 | **27** ± 0 | *n.s* |
| **HR** | *bpm* | **464** ± 9 | **442** ± 14 | *n.s* | **459** ± 27 | **448** ± 14 | *n.s* |
| **AWT ED** | *mm* | **0,8** ± 0,05 | **1,0** ± 0,02 | *p<0,01* | **0,8** ± 0,02 | **1,0** ± 0,02 | *p<0,001* |
| **AWT ES** | *mm* | **1,2** ± 0,05 | **1,5** ±0,05 | *p<0,01* | **1,3** ± 0,1 | **1,4** ±0,04 | *n.s* |
| **PWT ED** | *mm* | **0,8** ± 0,03 | **1,0** ±0,02 | *p<0,05* | **0,8** ± 0,02 | **1,0** ± 0,02 | *p<0,001* |
| **PWT ES** | *mm* | **1,1** ± 0,1 | **1,3** ± 0,03 | *p<0,05* | **1,1** ± 0,08 | **1,2** ± 0,04 | *n.s* |
| **LVEDD** | *mm* | **4,2** ± 0,1 | **4,3** ± 0,1 | *n.s* | **3,9** ± 0,04 | **4,7** ±0,1 | *p<0,001* |
| **LVESD** | *mm* | **2,9** ± 0,1 | **2,9** ± 0,1 | *p<0,05* | **2,2** ± 0,1 | **3,6** ±0,1 | *p<0,001* |
| **FS** | *%* | **30** ± 2 | **30** ± 1 | *n.s* | **40** ± 3 | **22** ± 1 | *p<0,001* |
| **VTI** | *cm* | **2,7** ± 0,1 | **2,6** ± 0,1 | *n.s* | **2,6** ± 0,1 | **2,7** ± 0,2 | *n.s* |
| **SV** | *µL* | **78** ± 4 | **64** ± 2 | *p<0,01* | **64** ± 4 | **62** ± 4 | *n.s* |
| **CO** | *mL/min* | **35** ± 2 | **30** ± 1 | *p<0,001* | **32** ± 3 | **27** ± 1 | *p<0,001* |
| **EDV** | *µL* | **65** ± 3 | **89** ±4 | *p<0,05* | **66** ± 3 | **109** ± 4 | *p<0,001* |
| **ESV** | *µL* | **30** ± 5 | **43** ± 3 | *n.s* | **25** ± 3 | **65** ± 3 | *p<0,001* |
| **EF** | *%* | **59** ± 4 | **53** ± 2 | *n.s* | **74** ± 2 | **44** ± 1 | *p<0,001* |

*BW*, body weight; *HR*, hear rate; *AWT ED*, anterior wall thickness end diastole; *AWT ES*, anterior wall thickness end systole; *PWT ED*, posterior wall thickness end diastole; *PWT ES*, posterior wall thickness end systole; *LVEDD*, left ventricular end diastolic diameter; *LVESD*, left ventricular end systolic diameter; *FS*, fractional shortening: *VTI*, velocity time integral; *SV*, stroke volume; *CO*, cardiac output; *EDV*, end diastolic volume; *ESV* end systolic volume; *EF*, ejection fraction. Two-way ANOVA, sham vs. TAC

Table S2. **Echocardiography female Balb/c TAC vs sham**

| | | **3 weeks** | | | **6 weeks** | | | **8 weeks** | | |
|---|---|---|---|---|---|---|---|---|---|---|
| **parameter** | | **sham** | **TAC** | ***vs sham*** | **sham** | **TAC** | ***vs sham*** | **sham** | **TAC** | ***vs sham*** |
| ***n*** | | ***23*** | ***17*** | | ***23*** | ***17*** | | ***8*** | ***10*** | |
| **BW** | g | **25** ± 0,3 | **24** ± 0,4 | *n.s* | **27** ± 0,3 | **25 ±** 0,3 | *n.s* | **27** ± 0,6 | **26** ± 0,3 | *n.s* |
| **HR** | *bpm* | **357** ± 10 | **324** ± 12 | *n.s* | **358** ± 9 | **381** ± 13 | *n.s* | **355** ± 21 | **329** ± 18 | *n.s* |
| **AWT ED** | *mm* | **0,8** ± 0,03 | **0,9** ± 0,04 | *n.s* | **0,8** ± 0,03 | **0,9** ± 0,04 | *n.s* | **0,8** ± 0,04 | **0,9** ± 0,04 | *n.s* |
| **AWT ES** | *mm* | **1,2** ± 0,04 | **1,2** ± 0,1 | *n.s* | **1,3** ± 0,04 | **1,2** ±0,1 | *n.s* | **1,3** ± 0,04 | **1,2** ±0,04 | *n.s* |
| **PWT ED** | *mm* | **0,9** ± 0,03 | **1,0** ± 0,05 | *n.s* | **0,9** ± 0,03 | **0,9** ± 0,04 | *n.s* | **1,0** ± 0,05 | **0,9** ± 0,05 | *n.s* |
| **PWT ES** | *mm* | **1,2** ± 0,04 | **1,2** ± 0,1 | *n.s* | **1,2** ± 0,1 | **1,1** ± 0,03 | *n.s* | **1,3** ± 0,1 | **1,1** ± 0,1 | *n.s* |
| **LVEDD** | *mm* | **3,7** ± 0,0 | **3,8** ± 0,1 | *n.s* | **3,7** ± 0,1 | **4,3** ± 0,1 | *p<0,001* | **3,7** ± 0,1 | **4,2** ± 0,1 | *p<0,001* |
| **LVESD** | *mm* | **2,5** ± 0,1 | **2,7** ± 0,1 | *n.s* | **2,5** ± 0,1 | **3,4** ± 0,2 | *p<0,001* | **2,4** ± 0,2 | **3,4** ± 0,1 | *p<0,001* |
| **FS** | *%* | **31** ± 2 | **28** ± 2 | *n.s* | **33** ± 2 | **22** ± 2 | *p<0,01* | **35** ± 3 | **18** ± 2 | *p<0,01* |
| **VTI** | *cm* | **2,6** ± 0,1 | **2,6** ± 0,1 | *n.s* | **2,5** ± 0,1 | **2,2** ± 0,1 | *n.s* | **2,5** ± 0,2 | **2,0** ± 0,2 | *n.s* |
| **SV** | *µL* | **47** ± 2 | **49** ± 3 | *n.s* | **51** ± 3 | **50** ± 4 | *n.s* | **47** ± 5 | **35** ± 3 | *p<0,05* |
| **CO** | *mL/min* | **18** ± 1 | **16** ± 1 | *n.s* | **18** ± 1 | **19** ± 2 | *n.s* | **17** ± 2 | **11** ± 1 | *p<0,05* |
| **EDV** | *µL* | **59** ± 2 | **62** ± 4 | *n.s* | **58** ± 3 | **83** ± 5 | *p<0,001* | **58** ± 3 | **79** ± 5 | *p<0,01* |
| **ESV** | *µL* | **24** ± 2 | **29** ± 4 | *n.s* | **24** ± 3 | **49** ± 5 | *p<0,01* | **21** ± 3 | **50** ± 5 | *p<0,001* |
| **EF** | *%* | **60** ± 2 | **54** ± 3 | *n.s* | **61** ± 3 | **43** ± 4 | *p<0,01* | **64** ± 4 | **37** ± 3 | *p<0,001* |

*BW*, body weight; *HR*, hear rate; *AWT ED*, anterior wall thickness end diastole; *AWT ES*, anterior wall thickness end systole; *PWT E*D, posterior wall thickness end diastole; *PWT ES*, posterior wall thickness end systole; *LVEDD*, left ventricular end diastolic diameter; *LVESD*, left ventricular end systolic diameter; *FS*, fractional shortening: *VTI*, velocity time integral; *SV*, stroke volume; *CO*, cardiac output; *EDV*, end diastolic volume; *ESV* end systolic volume; *EF*, ejection fraction. Two-way ANOVA, sham vs. TAC

Table S3. **Echocardiography female C57Bl/6J TAC vs sham**

| | | 3 weeks | | | 6 weeks | | | 8 weeks | | |
|---|---|---|---|---|---|---|---|---|---|---|
| **parameter** | | **sham** | **TAC** | ***vs sham*** | **sham** | **TAC** | ***vs sham*** | **sham** | **TAC** | ***vs sham*** |
| ***n*** | | ***22*** | ***19*** | | ***22*** | ***19*** | | ***8*** | ***10*** | |
| **BW** | g | **23** ± 0,3 | **23** ± 0,2 | *n.s* | **24** ± 0,3 | **25** ± 0,3 | *n.s* | **24** ± 0,5 | **25** ± 0,4 | *n.s* |
| **HR** | *bpm* | **399** ± 12 | **360** ± 16 | *n.s* | **410** ± 14 | **432** ± 14 | *n.s* | **445** ± 12 | **438** ± 10 | *n.s* |
| **AWT ED** | *mm* | **0,8** ± 0,03 | **1,0** ± 0,05 | *p<0,01* | **0,9** ± 0,02 | **1,1** ± 0,05 | *p<0,01* | **0,9** ± 0,03 | **1,3** ± 0,03 | *p<0,001* |
| **AWT ES** | *mm* | **1,2** ± 0,05 | **1,5** ± 0,1 | *p<0,01* | **1,4** ± 0,04 | **1,5** ± 0,1 | *n.s* | **1,4** ± 0,05 | **1,7** ± 0,03 | *p<0,05* |
| **PWT ED** | *mm* | **0,8** ± 0,03 | **1,0** ± 0,03 | *p<0,01* | **0,9** ± 0,03 | **1,2** ± 0,1 | *p<0,001* | **0,9** ± 0,05 | **1,3** ± 0,1 | *p<0,01* |
| **PWT ES** | *mm* | **1,1** ± 0,04 | **1,3** ± 0,05 | *p<0,01* | **1,1** ± 0,05 | **1,4** ± 0,1 | *p<0,01* | **1,1** ± 0,1 | **1,6** ± 0,1 | *p<0,01* |
| **LVEDD** | *mm* | **3,9** ± 0,1 | **3,7** ± 0,1 | *p<0,05* | **3,8**± 0,1 | **3,7** ± 0,1 | *n.s* | **3,6** ± 0,1 | **3,8** ± 0,1 | *n.s* |
| **LVESD** | *mm* | **2,8** ± 0,1 | **2,5** ± 0,1 | *p<0,05* | **2,7** ± 0,1 | **2,8** ± 0,1 | *n.s* | **2,5** ± 0,1 | **2,9** ± 0,1 | *n.s* |
| **FS** | *%* | **28** ± 1 | **32** ± 2 | *n.s* | **29** ± 2 | **26** ± 2 | *n.s* | **31** ± 2 | **25** ± 2 | *n.s* |
| **VTI** | *cm* | **2,4** ± 0,1 | **2,5** ± 0,1 | *n.s* | **2,5** ± 0,1 | **1,9** ± 0,1 | *p<0,05* | **2,6** ± 0,1 | **1,9** ± 0,1 | *p<0,01* |
| **SV** | *µL* | **46** ± 2 | **56** ± 3 | *p<0,05* | **46** ± 3 | **42** ± 4 | *n.s* | **47** ± 3 | **46** ± 3 | *n.s* |
| **CO** | *mL/min* | **18** ± 1 | **19** ± 1 | *n.s* | **19** ± 1 | **19** ± 1 | *n.s* | **21** ± 2 | **20** ± 1 | *n.s* |
| **EDV** | *µL* | **68** ± 2 | **57** ± 3 | *p<0,01* | **65** ± 3 | **59** ± 4 | *n.s* | **59** ± 3 | **62** ± 5 | *n.s* |
| **ESV** | *µL* | **31** ± 2 | **23** ± 2 | *p<0,01* | **30** ± 3 | **32** ± 3 | *n.s* | **24** ± 2 | **32** ± 4 | *n.s* |
| **EF** | *%* | **54** ± 2 | **60** ± 2 | *n.s* | **55** ± 2 | **50** ± 3 | *n.s* | **60** ± 2 | **49** ± 3 | *p<0,05* |

*BW*, body weight; *HR*, hear rate; *AWT ED*, anterior wall thickness end diastole; *AWT ES*, anterior wall thickness end systole; *PWT E*D, posterior wall thickness end diastole; *PWT ES*, posterior wall thickness end systole; *LVEDD*, left ventricular end diastolic diameter; *LVESD*, left ventricular end systolic diameter; *FS*, fractional shortening: *VTI*, velocity time integral; *SV*, stroke volume; *CO*, cardiac output; *EDV*, end diastolic volume; *ESV* end systolic volume; *EF*, ejection fraction. Two-way ANOVA, sham vs. TAC

Table S4. **Echocardiographic evaluation of cardiac function after inhibition of lymphangiogenesis**

| parameter | | 3 weeks TAC | 3 weeks anti-VEGFR3 TAC | 3 weeks *vs TAC* | 6 weeks TAC | 6 weeks anti-VEGFR3 TAC | 6 weeks *vs TAC* |
|---|---|---|---|---|---|---|---|
| *n* | | *13* | *9* | | *13* | *13* | |
| **BW** | g | **26** ± 0.3 | **25** ± 0.3 | *n.s* | **28** ± 0.3 | **27** ± 0.6 | *n.s* |
| **HR** | *bpm* | **409** ± 13 | **424** ± 9 | *n.s* | **418** ± 19 | **448** ± 16 | *n.s* |
| **AWT ED** | *mm* | **1,0** ± 0,03 | **0.9** ± 0,03 | *n.s* | **0,9** ± 0,02 | **0,9** ± 0,04 | *n.s* |
| **AWT ES** | *mm* | **1,5** ±0,05 | **1,4** ±0,02 | *n.s* | **1,4** ±0,04 | **1,4** ±0,04 | *n.s* |
| **PWT ED** | *mm* | **0,9** ±0,03 | **0,9** ± 0,03 | *n.s* | **0,9** ± 0,03 | 0.9 ± 0,03 | *n.s* |
| **PWT ES** | *mm* | **1,3** ± 0,03 | **1,3** ± 0.04 | *n.s* | **1,2** ± 0,04 | **1,2** ± 0,4 | *n.s* |
| **LVEDD** | *mm* | **4,2** ± 0,1 | **4,7** ± 0.1 | *p<0.05* | **4,8** ±0,1 | **4,6** ± 0.1 | *n.s* |
| **LVESD** | *mm* | **2,7** ± 0,1 | **3,4** ± 0,1 | *p<0.05* | **3,6** ±0,1 | **3,4** ± 0,2 | *n.s* |
| **FS** | *%* | **34** ± 2 | **28** ± 1 | *p<0.05* | **26** ± 1 | **27** ± 2 | *n.s* |
| **VTI** | *cm* | **2,6** ± 0,1 | **2,5** ± 0,1 | *n.s* | **2,8** ± 0,2 | **2,4** ± 0,2 | *p=0.1* |
| **SV** | *µL* | **75** ± 3 | **70** ± 6 | *n.s* | **78** ± 6 | **67** ± 5 | *p=0.1* |
| **CO** | *mL/min* | **30** ± 1 | **30** ± 1 | *n.s* | **32** ± 2 | **30** ± 2 | *n.s* |
| **EDV** | *µL* | **84** ± 5 | **97** ± 6 | *n.s* | **108**± 7 | **105** ± 6 | *n.s* |
| **ESV** | *µL* | **35** ± 3 | **46** ± 4 | *p<0.05* | **55** ± 5 | **53** ± 4 | *n.s* |
| **EF** | *%* | **61** ± 2 | **55** ± 2 | *n.s* | **50**± 2 | **51** ± 2 | *n.s* |

*BW*, body weight; *HR*, hear rate; *AWT ED*, anterior wall thickness end diastole; *AWT ES*, anterior wall thickness end systole; *PWT E*D, posterior wall thickness end diastole; *PWT ES*, posterior wall thickness end systole; *LVEDD*, left ventricular end diastolic diameter; *LVESD*, left ventricular end systolic diameter; *FS*, fractional shortening: *VTI*, velocity time integral; *SV*, stroke volume; *CO*, cardiac output; *EDV*, end diastolic volume; *ESV* end systolic volume; *EF*, ejection fraction. Two-way ANOVA, TAC vs. anti-VEGFR3-treated TAC

### Supplemental methods

#### Human samples

Five micrometer thick sections were prepared from paraffin-embedded human septal samples. Standard immunohistological protocols, including citrate antigen retrieval, were used to reveal podoplanin-positive lymphatic vessels using a mouse anti-human podoplanin/D2-40 antibody validated for clinical practice (Dako, # M3619, diluted *1:50*). Signal was revealed following incubation with HRP-conjugated donkey anti-mouse secondary, followed by DAB detection kit (Vector Laboratories, Burlingame, CA). All patient slides were processed together for uniformity. Lymphatics were examined using a Zeiss axiovision light microscope equipped with CCD cameras at *x10.* Lymphatic vessels, identified as Podoplanin$^+$ vascular structures in the subendocardium, were counted by an observer blinded to the patient category. Total lymphatic density, open lymphatic density (lumenized vessels), lymphatic vessel diameter, and density of perivascular lymphatics (defined as lymphatics within 200 µm distance of a large blood vessel) were determined. On average for each patient 1.2 ±0.2 mm² of the septal sample was analyzed to determine lymphatic vessel sizes and densities.

#### Experimental Model

Male and female C57Bl/6J and female Balb/c mice (22-24 g) were obtained from Janvier. Animal housing and experiments were in accordance with National Institutes of Health guidelines, and the study was ethically approved by the Normandy University regional review board according to French and EU legislation (01181.01 / APAFIS #8157-2016121311094625 v5 and APAFIS #23175-2019112214599474 v6).

Minimally-invasive transversal aortic banding constriction (TAC) was performed on 8 week-old adult male and female C57Bl/6J and female Balb/c mice, as previously described[1]. Mice were anaesthetized by intraperitoneal injection of ketamine (100 mg kg $^{-1}$ Imalgene®) and xylazine (10 mg kg $^{-1}$ Rompun® 2%, Bayer Health Care) and placed on mechanical ventilation. The operator performed a minimal thoracotomy with an incision at the level of the first intercostal space. The aortic arch was visualized under low-power magnification. A snare, made of 7-0 polypropylene suture, was passed under the aorta between the origin of the right innominate and left common carotid arteries. Two suture bands were placed side-by-side to create an elongated stenosis and prevent internalization of the suture, as described[2]. A bent 26-gauge needle was placed next to the aortic arch, and the sutures were snugly tied around the needle and the aorta. After banding, the needle was quickly removed. The skin was closed, and mice recovered on a warming pad until fully awake. The sham procedure was identical except that the aorta was not banded. Buprenorphine (50 µg/kg, Buprecare®, Axcience) was injected subcutaneously 6 hours after surgery and twice per day until 3 days post-operation.

#### Echocardiography

Transthoracic echocardiography measurements (Doppler and M-mode) were performed in animals anaesthetized with isoflurane (1-2%), using a Vevo 3100 Imaging System (FUJIFILM). The heart was analysed in the two-dimensional mode in parasternal short-axis views. LV fractional shortening and ejection fraction were calculated using M mode imaging. To evaluate heart rate (HR) and velocity time integral (VTI), a pulsed Doppler of the LV outflow tract was performed, allowing calculation of stroke volume ($SV = \pi \times$ LV outflow radius$^2 \times$ VTI) and cardiac output ($CO = SV \times HR$). LV outflow radius was measured for each animal. Aortic flow was measured using pulsed Doppler to verify the efficacy of transversal aortic constriction compared to sham mice. Echocardiographic analyses were performed with VevoLAB software.

#### Immunohistochemistry

Murine cardiac samples were sectioned into a central slice, which was snap-frozen. Cardiac sections were cut on a cryostat (8 µm thickness) and collected on SuperFrost plus glass slides. After fixation in acetone for 10 min, non-specific binding sites were blocked in 3% BSA in PBS, followed by Biotin-Avidin Blocking kit (Thermo Scientific) when streptavidin (SA)-conjugates were used to detect biotinylated secondary antibodies. Primary antibodies (*see table s5*), diluted in 1% BSA in PBS, were incubated on the sections at r.t. for 1h, followed by repeated washing in PBS and incubation with secondary antibodies (*see table s6*) for 30 minutes to 1h. Double or triple stainings were performed sequentially, and negative controls included omission of primary antibodies. Slides were mounted in Vectashield containing DAPI, and images were acquired using x20 or x40 objectives on a Zeiss epifluorescence microscope (AxioImager J1) equipped with an apotome and Zen 2012 software (Zeiss). Images were analyzed by an operator blinded to the treatment groups using Fiji imaging software (NIH).

**Table S5** – primary antibodies and reagents used for immunohistochemistry

| antigen | article | supplier | species reactivity | host | dilution | Conc (µg/mL) |
|---|---|---|---|---|---|---|
| **alpha SMA-FITC** | *F3777* | Sigma Aldrich | mouse | mouse | 1/100 | **28** |
| **CD3** | *A0452* | DAKO | human, mouse | rabbit | 1/100 | **6** |
| **CD4** | *13-0042-82* | eBioscience | mouse | rat | 1/200 | **2.5** |
| **CD8a** | *553028* | BD | mouse | rat | 1/500 | **1** |
| **CD31/PECAM** | *553371* | BD | mouse | rat | 1/100 | **0.6** |
| **CD68** | *14-0681-82* | eBioscience | mouse | rat | 1/800 | **5** |
| **F4/80** | *MCA497R* | Abd Serotec | mouse | rat | 1/200 | **5** |
| **iNOS** | *ab15323* | Abcam | mouse | rabbit | 1/200 | **2** |
| **LYVE-1** | *103-PA50* | Reliatech | mouse | rabbit | 1/1000 | **0.4** |
| **Biotinylated LYVE-1** | *103-PA50Bi* | Reliatech | mouse | rabbit | 1/500 | **0.4** |
| **CD206/MRC1** | *ab64693* | Abcam | mouse | rabbit | 1/2500 | **4** |
| **Podoplanin** | *14-5381-82* | eBioscience | mouse | hamster | 1/10 000 | **1** |
| **VEGF-C** | *ab9546* | Abcam | mouse | rabbit | 1/500 | **2** |
| **WGA** | *FP-CE8070* | Interchim | | | 1/100 | **1** |

**Table S6** – secondary antibodies & reagents used for immunohistochemistry

| antigen | Fluorochrome | article | supplier | Conc (µg/ mL) |
|---|---|---|---|---|
| **Donkey anti-Rat** | **AF488** | *712-545-153* | Jackson Immunoresearch | **3** |

| | | | | |
|---|---|---|---|---|
| **Donkey anti-Rat** | **Cy3** | *712-166-153* | Jackson Immunoresearch | **3** |
| **Goat anti-Rat** | **AF647** | *A-21247* | Thermo Fisher Scientific/ Invitrogen | **0.8** |
| **Donkey anti-Rabbit** | **Cy3** | *711-165-152* | Jackson Immunoresearch | **3** |
| **Donkey anti-Rabbit** | **AF647** | *711-605-152* | Jackson Immunoresearch | **1.5** |
| **Donkey anti-Goat** | **DYLIGHT 488** | *A50-201D2* | Interchim | **1.3** |
| **Donkey anti-Goat** | **DYLIGHT 550** | *A50-201D3* | Interchim | **1.3** |
| **Streptavidin** | **Fluoprobe 547** | *FP-CA5570* | Interchim fluoprobes | **0.7** |
| **Streptavidin** | **Fluoprobe 647** | *FP-CA5640* | Interchim fluoprobes | **0.7** |
| **Goat anti-Hamster** | **AF488** | *A-21110* | Thermo Fisher Scientific / Invitrogen | **0.8** |

#### Image analysis of cardiac sections

Murine lymphatic vessels were defined as strongly Lyve1-positive structures, lacking macrophage markers (F4/80 or CD68), and positive for Podoplanin. Lymphatics are preferentially located in the subepicardium in rodent hearts. These Lyve1-positive, F4/80 or CD68-negative vessels clearly differ from $CD68^+$ or $F4\text{-}80^+$ macrophages, which either lack or display weaker Lyve1 expression. These macrophages, also differing in size and morphology from the elongated cell body and nucleus typical of lymphatic endothelial cells, were readily excluded from lymphatic counts. Photos were captured at *x20* or *x40*.

Due to the absence of conclusive markers to distinguish precollectors from capillaries in murine cardiac lymphatics, we have made use of the fact that precollectors preferentially run in a base-to-apex fashion, whereas lymphatic capillaries lack such consistent spatial organization. Hence, by evaluating vessels with a lumen parallel to horizontal cardiac sections, the precollector population of vessels is more frequently detected as vessels running perpendicular to the section (= open lumen vessels). To assess the size and frequency of open lymphatic vessels (diameter > 5 µm; a population enriched in precollector vessels), between 10-20 images from the LV free-wall were collected and analyzed for each mouse. The density (open vessels/$mm^2$) and lymphatic lumen sizes were measured and used to calculate mean vessel diameter. The parameter % open lymphatic area was calculated as the sum total of all measured lymphatic diameters divided by the total cardiac area analyzed for each mouse heart. On average for each mouse $2.17 \pm 0.25$ $mm^2$ of the LV was analyzed to determine lymphatic vessel sizes and % open lymphatic area.

Blood vessels were defined as strongly CD31-positive structures. These clearly differed from lymphatic vessels that displayed weaker CD31 signal. Blood to cardiomyocyte ratios were evaluated in sections double stained for CD31 and WGA imaged at *x40*. Individual endothelial cells were not counted, rather vessels structures were assessed. If two vessels were adjacent and partly overlapping, they were counted as two vessels. For one vessel that divides into two branches, two vessels were counted.

Arterioles were defined as smooth muscle α-actin-positive vessels. Images were captured at *x20*. Cardiac macrophages were defined as CD68 or F4/80 positive cells. M2-like macrophages were defined as $CD206^+$ cells, and M1 type macrophages as $iNOS^+$ $CD206^-$

cells. Total T cells were defined as CD3 positive cells, with subpopulations identified as CD8a$^+$ or CD4$^+$ cells. B cells were defined as B220$^+$ immune cells. Images were captured at *x20*.

Interstitial fibrosis was evaluated as cardiac area positive for either collagen I or collagen III in images captured at *x20*. Images were analyzed by an operator blinded to the treatment groups using Fiji imaging software (NIH).

##### Histology

Cardiac cryosections (8 µm) of mouse hearts were processed for Sirius Red staining as described[3]. Cardiac interstitial collagen density was evaluated in sections imaged on a light microscope (Zeiss) equipped with an *x40* objective. Perivascular fibrosis was similarly evaluated in images taken at *x20* on a Zeiss epifluorescence microscope (AxioImager J1) equipped with an apotome and Zen 2012 software (Zeiss). Images were analyzed by an operator blinded to the treatment groups using Fiji imaging software (NIH). Perivascular fibrosis is expressed as fibrotic area (µm²) surrounding arteries in the size-range of 5-50 µm in diameter.

##### Real-Time Polymerase Chain Reaction

Murine cardiac samples were collected at 3 and 8 weeks post-TAC. RNA extraction was performed with Trizol method. Briefly, samples were homogenized (45 sec.) in TRIZOL using Precellys tubes with beads. RNA was obtained after incubation, successively, in chloroform and isopropanol and three washes in ethanol. RNA quality was analyzed by Nanodrop 2000 (Thermo Fisher Scientific). cDNAs were generated using a reverse transcriptase after DNAse treatment. Real-time polymerase chain reaction was performed on a LightCycler 480 (Roche Molecular Biochemicals) with a commercially available mix (FastStart DNA Master SYBR Green I kit; Roche) on white 96-well plates. The reactions were performed in duplicate for each sample. Concentrations were calculated using a standard curve, obtained by serial dilutions of samples from an organ with high expression of the gene of interest. To normalize gene expression, values are expressed as ratios of a reference gene (β2m). Healthy sham animal ratios are then set as 100%, and values in TAC-operated animals are expressed relative to their respective sham controls. The sequences of the specific primers are detailed in table s7.

**Table S7** – RT-PCR primers used for cardiac expression analyses in mouse

| gene | sense | antisense | amplicon (bp) |
|---|---|---|---|
| ***B2m*** | GCTGCTACTCGGCGCTTCA | GCAGGCGTATGTATCAGTCTCAGT | 343 |
| ***Ccl2*** | CCCAATGAGTAGGCTGGAGA | GCTGAAGACCTTAGGGCAGA | 210 |
| ***Ccl21*** | TCCCTACAGTATTGTCCGAGGC | ATCAGGTTCTGCACCCAGCCTT | 141 |
| ***Flt4 (Vegfr3)*** | AGACTGGAAGGAGGTGACCACT | CTGACACATTGGCATCCTGGATC | 128 |
| ***Lyve1*** | ACCAGGTAGAGTCAGCGCAGAA | CAGGACACCTTTGCCATTCTTCC | 128 |
| ***Pdpn*** | ACAACCACAGGTGCTACTGGAG | GTTGCTGAGGTGGACAGTTCCT | 116 |
| ***Vegfa*** | CTGCTGTAACGATGAAGCCCTG | GCTGTAGGAAGCTCATCTCTCC | 119 |
| ***Vegfc*** | AGCCCACCCTCAATACCAG | GCTGCTCCAAACTCCTTCC | 154 |
| ***Vegfd*** | CTCCACCAGATTTGCGGCAACT | ACTGGCGACTTCTACGCATGTC | 112 |

| | | | |
|---|---|---|---|
| ***Nppa*** | TACAGTGCGGTGTCCAACACAG | TGCTTCCTCAGTCTGCTCACTC | 126 |
| ***Nppb*** | TCCTAGCCAGTCTCCAGAGCAA | GGTCCTTCAAGAGCTGTCTCTG | 98 |
| ***Il1β*** | TGGACCTTCCAGGATGAGGACA | GTTCATCTCGGAGCCTGTAGTG | 148 |
| ***Il6*** | TACCACTTCACAAGTCGGAGGC | CTGCAAGTGCATCATCGTTGTTC | 116 |
| ***TNFα*** | GGTGCCTATGTCTCAGCCTCTT | GCCATAGAACTGATGAGAGGGAG | 139 |

#### Whole mount

Prior to sacrifice, deeply anesthetized mice were perfused with warm saline solution, followed by perfusion-fixation with warm 3% paraformaldehyde (PFA). Hearts were removed and post-fixed (3% PFA) for 6h. Following dehydration in graded methanol baths, and post-fixation in Dent's fixative, samples were bleached in graded $H_2O_2$ baths. After extensive blocking of non-specific binding sites and tissue permeation with Triton-X100, cardiac lymphatics were visualized using antibodies reactive against Lyve1 (rabbit polyclonal, Reliatech, *1/500*), followed by a fluorescence-coupled secondary antibody (Cy5-conjugated donkey anti-rabbit, *1/500)*. Arterial vasculature was visualized using Cy3-conjugated anti-alpha-smooth muscle actin antibodies (mouse monoclonal, 1/500, Sigma-Aldrich). Extensive washing was performed to remove nonspecific binding. Hearts were clarified using a modified iDISCO+ protocol, as described[4], based on incubation in graded methanol baths followed by incubation in dichloromethane (DCM, Sigma-Aldrich, *#270997-12X100ML*) and dibenzyl ether (DBE, Sigma-Aldrich, *#108014-1KG*) before light sheet and confocal imaging.

#### *3D Imaging and Image Processing*

3D imaging was performed by light sheet microscopy as described[4] Acquisitions were performed with an ultramicroscope II (LaVision BioTec) using the ImspectorPro software (LaVision BioTec). The light sheet was generated by a laser (wavelengths 561 or 640 nm, Coherent Sapphire Laser, LaVision BioTec) focused using two cylindrical lenses. A binocular stereomicroscope (MXV10, Olympus) with an *x*2 objective (MVPLAPO, Olympus) was used at different magnifications (*x*0.8 and *x*4). A dipping cap protective lens, including correction optics for MVPLAPO x2 objective, was applied for working distances inferior or equal to 5.7mm. The corresponding zoom factors and numerical apertures are available at:
https://www.miltenyibiotec.com/FR-en/products/ultramicroscope-ii.html#scrollto_specs

Samples were placed in an imaging reservoir made of 100% quartz (LaVision BioTec) filled with DBE and illuminated from the side by the laser light sheet. A PCO Edge SCMOS CCD camera (2560× 2160 pixel size, LaVision BioTec) was used to capture images. The step size between each image was fixed at 6 µm (*x*0.8 zoom) or 2 µm (*x*3.2 zoom). All tiff images were generated in 16-bit.

For confocal laser scanning microscopy, imaging was performed with an upright fixed-stage TCS SP8 confocal microscope (Leica Microsystems, France) equipped with multiple laser lines (wavelengths 561 or 640 nm). In order to acquire deep confocal views, a *x25* objective was used (numerical aperture 0.95, working distance 2500 µm, water immersion). Images were captured using a hydrid detector (Hamamatsu) in photon counting z-stack mode.

##### Image Processing

Images, 3D volume, and movies were generated using Imaris x64 software (version 8.0.1, Bitplane). Z stack light sheet images were first converted to imaris file (.ims) using ImarisFileConverter and 3D reconstruction was performed using the "volume rendering" function. To facilitate image processing, images were converted to 8-bit format. Optical slices were obtained using the "orthoslicer" tool. 3D pictures and movies were generated using the "snapshot" and "animation" tools. Movie reconstruction with .tiff series were performed with NIH ImageJ (1.50e, Java 1.8.0_60, 64-bit).

Alternatively, 3D reconstruction and segmentation of cardiac lymphatic network in ultramicroscopy stacks were performed using custom software leveraging the Voreen framework analysis as previously described[5]. Briefly, the analysis workflow consists of the three basic steps: segmentation, skeletonization, and feature extraction. For segmentation of the lymphatic vessels, an interactive semiautomatic random walker segmentation was applied to a downsampled version of the Lyve1-positive channel.

Z stack confocal raw data images were deconvoluted through image processing with Huygens professional software (SVI, The Netherlands). Maximal intensity projection views were generated with Fiji Image J software (1.51u, Java 1.8.0_66, 64-bit).

#### Flow Cytometry

Animals were sacrificed under deep anaethezia (thiopenthal) and following cardiac perfusion with warmed physiological saline through the abdominal aorta, hearts were recovered. Left ventricular samples were minced with scalpels and placed in ice-cold DMEM medium. Single cell suspensions were prepared, as described[4], by incubation in tissue-dissociating solution (125 U/mL collagenase type XI, *#C7657*, Sigma; 450 U/mL collagenase type I, *#C0130*, Sigma; 60 U/mL hyaluronidase type I, *#H3506*, Sigma; and 60 U/mL DNase 1 *#DN25*, Sigma) coupled to gentleMACS Dissociator (MACS; Miltenyi Biotec, Auburn, CA). Digested tissues were washed with DMEM culture medium and filtered to remove undigested tissue pieces (80 µm mesh, and 40 µm mesh, BD Biosciences).

Single cell suspensions were prepared in FACS buffer (1% BSA in PBS with 3 mM EDTA). To block nonspecific binding of antibodies to Fcγ receptors, isolated cells were first incubated with Fc-Block for 15 minutes at 4°C. Subsequently, cells were stained with specific antibody cocktails (see *table s8*) for 20 minutes at r.t, followed by washing with FACS buffer and resuspension in 600 µL FACS buffer. To quantify immune cells per mg heart, fluobeads (Flow-count Fluorospheres, Beckman coulter *#7547053*) were added to each sample prior to analysis. Monocytes were defined as live $CD45^+/CD11b^+/SSC^{low}/CD11c^-$ or $MHC^{IIlow}$ cells, whereas macrophages were identified as $F4/80^+$ positive subset. T cells were identified as live $CD45^+/CD11b^-/CD3^+$ cells. B cells were identified as live $CD45^+/CD19^+$ or $B220^+$ cells. NK cells were defined as live $CD45^+/CD3^-/NK1.1^+$ cells. Flow cytometry was performed on an 18-color **LSRFortessa** (BD Biosciences) followed by analysis using FlowJo software (TreeStar, Inc, San Carlos, CA). Results are expressed as immune cells per mg heart tissue.

**Table S8** – primary antibodies used for flow cytometry analyses in mouse

| Antigen | Fluorochrome | Source & catalogue # |
|---|---|---|
| **CD16/32 (Fc block)** | - | BD Pharmingen 553142 |
| **Fixable Viability Dye (L/D)** | **eFluor™ 455UV** | ebioscience 65-0868-14 |
| **CD45** | **PerCP** | Sony Biotechnology 1115650 |

| | | |
|---|---|---|
| **Ly-6C** | **APC/Cy7** | Sony Biotechnology 1240130 |
| **LY6G** | **PerCP/Cy5.5** | Biolegend clone 1A8 |
| **CD11b** | **FITC** | BD Pharmingen 553310 |
| **CD11b** | **Pacific blue** | Biolegend clone M1/70 |
| **CD11c** | **PE-Dazzle** | Sony Biotechnology 1186740 |
| **F4/80** | **BV605** | Sony Biotechnology 1215665 |
| **CD64** | **PE/Cy7** | Biolegend clone X54-5/7.1 |
| **MHCII (I-A/I-E)** | **AF700** | Sony Biotechnology 1138110 |
| **CD206** | **APC** | Sony Biotechnology 1308540 |
| **CD206** | **BV605** | Biolegend clone MCA2235F |
| **CD192 (CCR2)** | **PE/Cy7** | Sony Biotechnology 1353060 |
| **CX3CR1** | **BV421** | Sony Biotechnology 1345115 |
| **CD172a** | **APC** | Sony Biotechnology 1320070 |
| **CD45R (B220)** | **PE** | BD Pharmingen 553090 |
| **CD19** | **BV711** | Biolegend 115555 |
| **CD3e** | **APC** | BD Pharmingen 553066 |
| **CD3** | **BV650** | Biolegend 100229 |
| **CD4** | **APC/Cy7** | Sony Biotechnology 1102070 |
| **CD8a** | **AF700** | Sony Biotechnology 103650 |
| **FoxP3** | **AF488** | Sony Biotechnology 2200055 |
| **NK1-1** | **PE/Cy7** | BD Pharmingen 552878 |

#### References


1. Hu P, Zhang D, Swenson L, Chakrabarti G, Abel ED, Litwin SE. Minimally invasive aortic banding in mice: effects of altered cardiomyocyte insulin signaling during pressure overload. *American Journal of Physiology - Heart and Circulatory Physiology*. 2003;285:H1261–H1269.

2. Lygate CA, Schneider JE, Hulbert K, Hove M, Sebag-Montefiore LM, Cassidy PJ, Clarke K, Neubauer S. Serial high resolution 3D-MRI after aortic banding in mice: band internalization is a source of variability in the hypertrophic response. *Basic Research in Cardiology*. 2006;101:8–16.

3. Henri O, Pouehe C, Houssari M, Galas L, Nicol L, Edwards-Lévy F, Henry J-P, Dumesnil A, Boukhalfa I, Banquet S, Schapman D, Thuillez C, Richard V, Mulder P, Brakenhielm E. Selective Stimulation of Cardiac Lymphangiogenesis Reduces Myocardial Edema and Fibrosis Leading to Improved Cardiac Function Following Myocardial Infarction. *Circulation*. 2016;133:1484–1497; discussion 1497.

4. Houssari M, Dumesnil A, Tardif V, Kivelä R, Pizzinat N, Boukhalfa I, Godefroy D, Schapman D, Hemanthakumar KA, Bizou M, Henry J-P, Renet S, Riou G, Rondeaux J, Anouar Y, Adriouch S, Fraineau S, Alitalo K, Richard V, Mulder P, Brakenhielm E. Lymphatic and Immune Cell Cross-Talk Regulates Cardiac Recovery After Experimental Myocardial Infarction. *Arterioscler Thromb Vasc Biol*. 2020;40:1722–1737.

5. Hägerling R, Drees D, Scherzinger A, Dierkes C, Martin-Almedina S, Butz S, Gordon K, Schäfers M, Hinrichs K, Ostergaard P, Vestweber D, Goerge T, Mansour S, Jiang X, Mortimer PS, Kiefer F. VIPAR, a quantitative approach to 3D histopathology applied to lymphatic malformations. *JCI Insight*. 2017;2.
